# A molecular integrator of sleep duration and interruption

**DOI:** 10.64898/2026.07.03.736427

**Authors:** Elizabeth I. Tilden, Antonio J. Fontenele, Kane M. Goggans, Sophie Ma, Drake Gorecki, Zachary D. Berriman-Rozen, Anna Oldenborg, Woodrow L. Shew, Yao Chen

## Abstract

Sleep is regulated across multiple timescales. Transitions between sleep and wake happen within seconds^1,2^; individual sleep bouts last minutes to hours^3^; and homeostatic sleep need has classically been tracked across multiple bouts^4^. Rapid sleep-to-wake transitions are driven by identified neurons, circuits, and neuromodulators^1^, while slow wave activity correlates with sleep need across hours^5,6^. However, no signal has been shown to encode sleep history within individual sleep bouts, the timescale at which the brain must continuously monitor how much sleep has occurred and how likely waking is at any given moment. Biochemical signals downstream of sleep/wake-associated neuromodulators display slower dynamics than the neuromodulators themselves^7,8^, making them candidate encoders of within-bout sleep history. Here, by measuring protein kinase A substrate phosphorylation (PKA-SP) in real time in freely behaving mice^7–9^, we show that membrane PKA-SP decreases exponentially within each sleep bout with consistent kinetics across bouts, integrates sleep duration and sleep interruption, and continuously forecasts moment-to-moment waking probability. Following sleep deprivation, PKA-SP reaches lower levels at the end of sleep bouts, correlating with increased sleep need dissipation. These findings identify a molecular signal encoding within-bout sleep history, revealing how biochemical dynamics bridge fast arousal circuits and the slow timescale of classical sleep homeostasis.

## Introduction

Sleep is critical for survival and health^10–19^, and although we spend one third of our lives sleeping, the brain mechanisms that coordinate when and for how long we sleep remain incompletely understood. Sleep is organized into bouts of variable durations^3^. While significant progress has been made in identifying neurons and circuits that promote rapid transitions between sleep and wake^1,2,20–26^, what accounts for the timing of these transitions remains unclear. To date, no signal has been shown to track the passage of time within a sleep bout, largely because the tools to measure signals at within-bout temporal resolution did not exist until recently^7,8,27,28^. Prior candidate signals occupy two extremes of temporal dynamics: slow wave activity (SWA) declines across bouts but remains largely flat within individual bouts^5,29,30^, whereas neuromodulators such as adenosine display rapid transitions at sleep-wake boundaries^31,32^ rather than the gradual dynamics expected for a signal encoding sleep history within a bout. Because neuromodulators converge onto biochemical signaling pathways^33,34^, and these signals often display slower kinetics than the neuromodulators themselves^7,8^, biochemical signals such as kinases and second messengers are candidate molecular encoders of gradual within-bout sleep history.

We hypothesize that specific molecular signals dynamically encode sleep history throughout a sleep bout, tracking the moment-to-moment probability of waking. This hypothesis is inspired by the sleep need concept, stemming from the two-process model, which proposed that homeostatic sleep need dissipates during sleep in a time-dependent manner^4,35,36^. Classically, this dissipation is conceptualized across hours and multiple bouts, with SWA as the best available marker^5^. However, if sleep need dissipates gradually during sleep, this dissipation must begin within individual bouts, because each bout is a fundamental unit of sleep and because even brief naps can partially relieve sleep need^37,38^. This reasoning predicts the existence of molecular signals that change gradually within sleep bouts and encode the cumulative history of sleep within each bout. Whether such signals exist, what properties they encode, and how they relate to sleep need dissipation remain open questions.

If molecular signals encode sleep history within bouts, they would be expected to (1) change gradually across the duration of each sleep bout; (2) integrate sleep duration and sleep interruption into a cumulative representation of sleep history; (3) track the progressive increase in waking probability across the sleep bout; and (4) show greater change per bout when sleep need is higher, for example after sleep deprivation. Beyond these primary criteria, such a signal would also be expected to exhibit consistent dynamics across brain regions, reflecting the global nature of sleep regulation^39–41^, and to be influenced by neuromodulators differentially released between sleep and wake^1,31,32,42–45^.

Protein kinase A (PKA) is a molecular integrator of many of these sleep/wake-associated neuromodulators^34,46–49^. Importantly, PKA substrate phosphorylation (PKA-SP) dynamics change on the timescale of tens of seconds to minutes in response to neuromodulators^7,8^, positioning PKA-SP as a temporal integrator capable of capturing the cumulative neuromodulatory history within a sleep bout. Intriguingly, in conditions where sleep need is high, such as in *Sleepy* mouse models and wild type sleep-deprived mice, phosphoproteome analyses of brain lysates identified increased phosphorylation of protein substrates by five kinases, including PKA^39^. These phosphoproteome findings are consistent with additional evidence showing that the phosphorylation-dephosphorylation balance of PKA and its cognate phosphatases regulates sleep-wake behaviors^50–52^. Furthermore, PKA phosphorylates substrates important for synaptic scaling and sleep need^39,53–59^.

However, existing evidence implicating PKA-SP in sleep need is both indirect and difficult to reconcile across studies. The reliance on static biochemical snapshots from mixed cell populations has yielded apparently contradictory results. Phosphoproteome studies reported increased PKA-SP following sleep deprivation, consistent with a role in sleep need accumulation^39^. Other biochemical methods found decreased cyclic AMP (cAMP) and PKA-SP in response to the same manipulation, which is consistent with a role for PKA-SP in sleep-dependent memory consolidation^60^ where increased PKA activity during sleep supports synaptic plasticity^61^. Thus, it is unclear from the existing studies whether increased sleep pressure correlates with increased or decreased PKA-SP. These discrepancies might reflect the challenge of capturing highly dynamic, cell-type specific signals with population-level static assays and raise the possibility that PKA-SP dynamics serve multiple functions during sleep. Techniques with higher spatiotemporal resolution are critical to addressing these challenges.

Here we probe PKA-SP dynamics by recording PKA-SP in real time across sleep-wake cycles in freely behaving mice. We reveal striking subcellular differences between intracellular and membrane PKA-SP dynamics. Membrane PKA-SP decreases exponentially during each sleep bout, with consistent parameters across bouts. By dissipating during sleep and rising during microarousals, membrane PKA-SP integrates both sleep duration and sleep interruption to predict waking probability. Further, membrane PKA-SP dissipates to a lower level at the end of each sleep bout in sleep-deprived animals. Additionally, membrane PKA-SP displays similar dynamics across cortical and hippocampal regions potentially reflecting the global nature of sleep regulation. Membrane PKA-SP is also regulated by neuromodulators implicated in sleep need. Together, these findings provide the first direct evidence of a molecular signal whose in vivo dynamics track waking probability within sleep bouts, consistent with a signal that computes the relative dissipation of sleep need at the bout level. These results reveal how a subcellular signal functions as both a timer of sleep duration and a reporter of sleep interruption, encoding cumulative sleep history within each bout.

## Results

### Compartment-specific PKA-SP dynamics across sleep-wake cycles

To test whether PKA-SP dynamics fulfill the predicted criteria for a within-bout signal of sleep history, we used fluorescence lifetime photometry (FLiP)^9^ with a fluorescence lifetime-compatible A kinase activity reporter (FLIM-AKAR) that we previously developed^7,8^. Upon phosphorylation, the consensus substrate region of FLIM-AKAR interacts with the phosphopeptide binding domain, bringing the donor and acceptor fluorophore closer and resulting in Förster resonance energy transfer (FRET)^8^. Using FLiP, we can measure the binding fraction of FLIM-AKAR with 1 Hz resolution, representing real-time PKA-SP resulting from the combined actions of PKA and phosphatase activity. Since PKA phosphorylates different protein substrates with distinct cellular functions in different subcellular compartments, we used either the freely diffusible FLIM-AKAR to record PKA-SP in the intracellular compartment, or FLIM-AKAR with a farnesylation domain (FLIM-AKAR_m_) to record PKA-SP at the plasma membrane.

We delivered adeno-associated virus (AAV) encoding Cre-dependent FLIM-AKAR targeted to either compartment in excitatory neurons in the dorsal CA1 region of the hippocampus of *Tg(CamKIIα-Cre)* mice^62^ and implanted an optical fiber at the same site (**Fig. 1a**). As expected, FLIM-AKAR was expressed highly inside the cell, clearly visible in the soma and proximal dendrites (**Fig. 1b**). FLIM-AKAR_m_ was enriched in the membrane of soma and neuronal processes, including dendritic spines (**Fig. 1b**). We performed simultaneous FLiP (1Hz), electroencephalogram (EEG; 400Hz), electromyography (EMG; 400Hz), and video recordings (25 frames per second) in freely behaving mice for 24-48 hours. Using a custom sleep scoring program^63^, we identified non-rapid eye movement (NREM) sleep, rapid eye movement (REM) sleep, microarousals, and wake stages (**Fig. 1c**).

**Figure 1.**
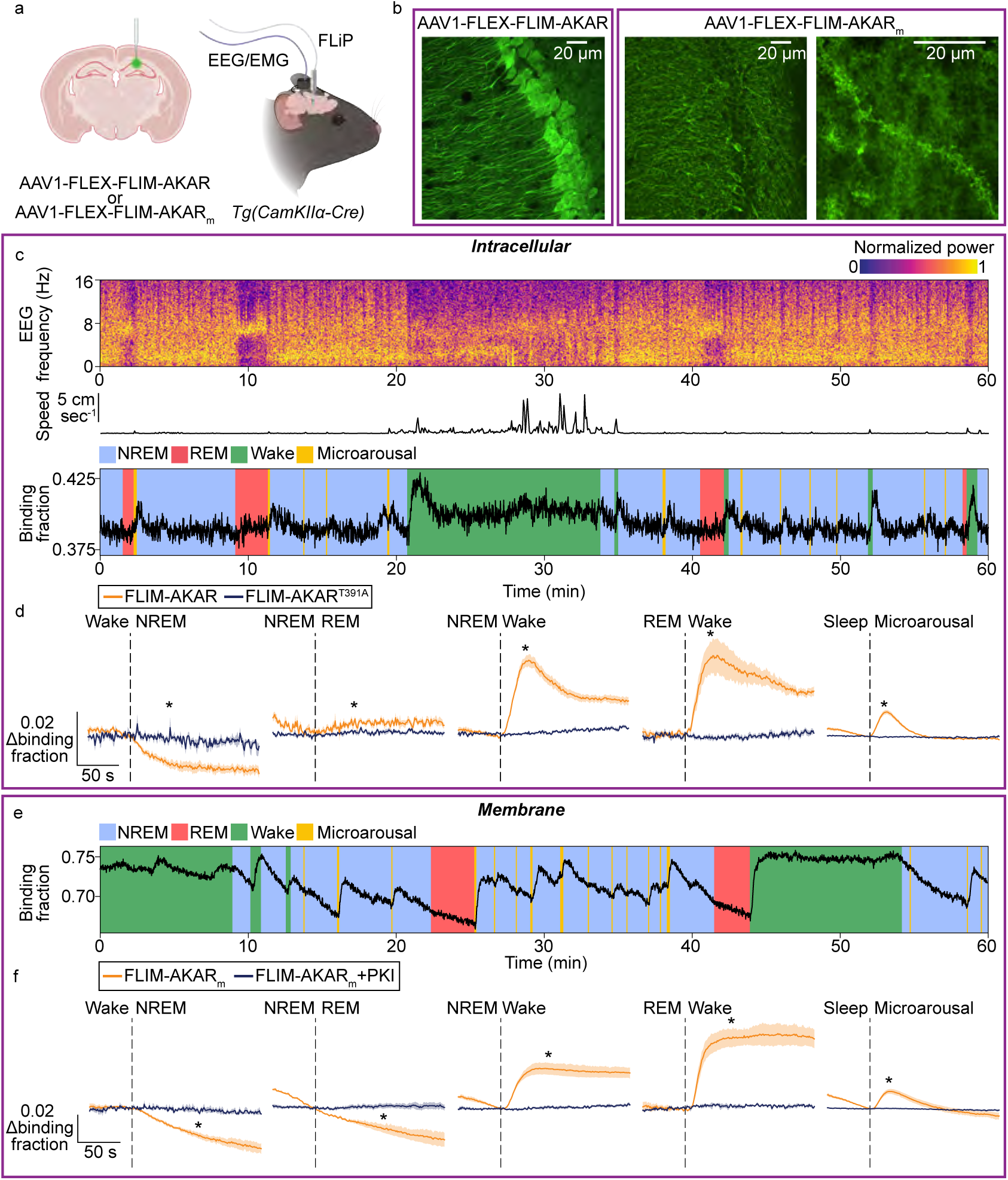
PKA substrate phosphorylation shows compartment-specific dynamics associated with sleep-wake states. a, Schematic illustrating that AAV1-FLEX-FLIM-AKAR or AAV1-FLEX-FLIM-AKARm was delivered to excitatory neurons of hippocampal CA1 for combined electroencephalogram (EEG)/electromyography (EMG) and fluorescence lifetime photometry (FLiP) recordings in Tg(CamKIIα-Cre) mice. b, Confocal images showing subcellular sensor localization of intracellular (left: optical section) and membrane-targeted (middle: optical section, right: max projection, 15 slices) FLIM-AKAR sensors. High resolution image (right) allows for visualization of spines. c, Example EEG spectrogram, animal speed, and intracellular FLIM-AKAR recordings with sleep stages over an hour. d, Behavior state transition-triggered change in the binding fractions of FLIM-AKAR (n=5 mice) or the non-phosphorylatable point mutant FLIM-AKART391A (n=4 mice). Data show mean ± standard error of the mean (SEM) (Welch’s t-test; FLIM-AKAR vs. FLIM-AKART391A). e, Example recording from the membrane-targeted reporter FLIM-AKARm with sleep stages over one hour. f Behavior state transition-triggered averages of the binding fraction of the membrane-targeted reporter FLIM-AKARm in the presence (n=4 mice) or absence (n=4 mice) of PKI. Data show mean ± SEM (Welch’s t-test; FLIM-AKARm vs. FLIM-AKARm + PKI). *: p<0.05. See details in Extended Data Table 1.

For intracellular PKA-SP, our recordings revealed that FLIM-AKAR binding fraction transiently increased upon waking, followed by partial recovery to an intermediate level. Upon entry into NREM, FLIM-AKAR binding fraction rapidly reduced to a plateau within a minute of NREM onset (**Fig. 1c, d)**. Microarousals resulted in a transient increase in FLIM-AKAR binding fraction. These dynamics are consistent with a recent report of cAMP dynamics across sleep-wake cycles^64^. These observations are robust across multiple sleep-wake bouts within one mouse (**ED Fig. 1a**), and across mice (**Fig. 1d).** These dynamics were not observed with a non-phosphorylatable mutant FLIM-AKAR (FLIM-AKAR^T391A^)^8^, where the threonine phosphorylation site in the PKA substrate consensus motif is replaced with an alanine **(Fig. 1d)**.

Remarkably, PKA-SP displayed distinct sleep-associated dynamics at the membrane compared to the intracellular compartment (**Fig. 1e, f**). Upon waking, FLIM-AKAR_m_ binding fraction rapidly increased to a sustained high level. It decreased gradually during both NREM and REM sleep, interrupted by transient rises following microarousals (**Fig. 1e, f)**. The dynamics showed consistency across transitions within one animal (**ED Fig. 1b)** and across all animals (**Fig. 1f**). Additionally, FLIM-AKAR_m_ dynamics were abolished in the presence of the genetically-encoded, AAV-delivered PKA inhibitory peptide (PKI) (**Fig. 1f**), which inhibits PKA with high specificity^8^, indicating the sleep-wake-associated changes of FLIM-AKAR_m_ specifically reflect membrane PKA-SP. Importantly, to test whether the asymptotic binding fraction during wake was due to a ceiling effect of the sensor, we expressed and stimulated photoactivatable adenylyl cyclase (biPAC)^65^ while performing simultaneous FLiP recording and found that FLIM-AKAR_m_ binding fraction could increase above wake levels (**ED Fig. 1c, d).** Similarly, the slow dissipation of PKA-SP during sleep is not limited by sensor kinetics, as a faster decrease in the binding fraction of FLIM-AKAR_m_ was observed in response to a muscarinic acetylcholine receptor antagonist (**ED Fig. 1e, f**). Thus, membrane and intracellular PKA-SP exhibit distinct dynamics across sleep-wake cycles, suggesting that the same molecular signal encodes different information depending on its subcellular location.

Notably, the gradual decrease in membrane PKA-SP throughout the duration of a sleep bout and its transient increases from microarousals suggest that PKA-SP could integrate both sleep duration and sleep interruption to track proximal sleep history. Furthermore, its gradual dissipation during sleep and lack of accumulation during wake suggest it does not encode absolute sleep need and instead matches the predicted dynamics of a signal encoding the bout-to-bout dissipation of sleep need. Unless otherwise noted, all subsequent references to PKA-SP refer specifically to the membrane-compartment signal as measured by FLIM-AKAR_m_.

### Membrane PKA-SP dynamics are consistent across cortico-hippocampal regions

The within-bout dynamics of membrane PKA-SP raises the question of whether these properties are specific to the hippocampal circuit or a general feature of membrane PKA-SP in excitatory neurons, as would be expected for a signal tracking global sleep history^39–41^. To address this, we recorded membrane PKA-SP dynamics in excitatory neurons across cortical regions with distinct functional roles: visual cortex (V1), motor cortex (M1), and medial prefrontal cortex (mPFC), a region implicated in regulating sleep need^66^. Membrane PKA-SP in these three brain regions exhibited dynamics similar to those in hippocampal CA1 regions: PKA-SP was high during wake, gradually declined during sleep, and transiently increased during microarousals, with slightly different slopes during REM bouts (**Fig. 2, ED Fig. 2**). Thus, the within-bout dynamics of membrane PKA-SP are consistent across cortical and hippocampal regions, suggesting that these properties reflect a general feature of excitatory neurons rather than a circuit-specific phenomenon.

**Figure 2.**
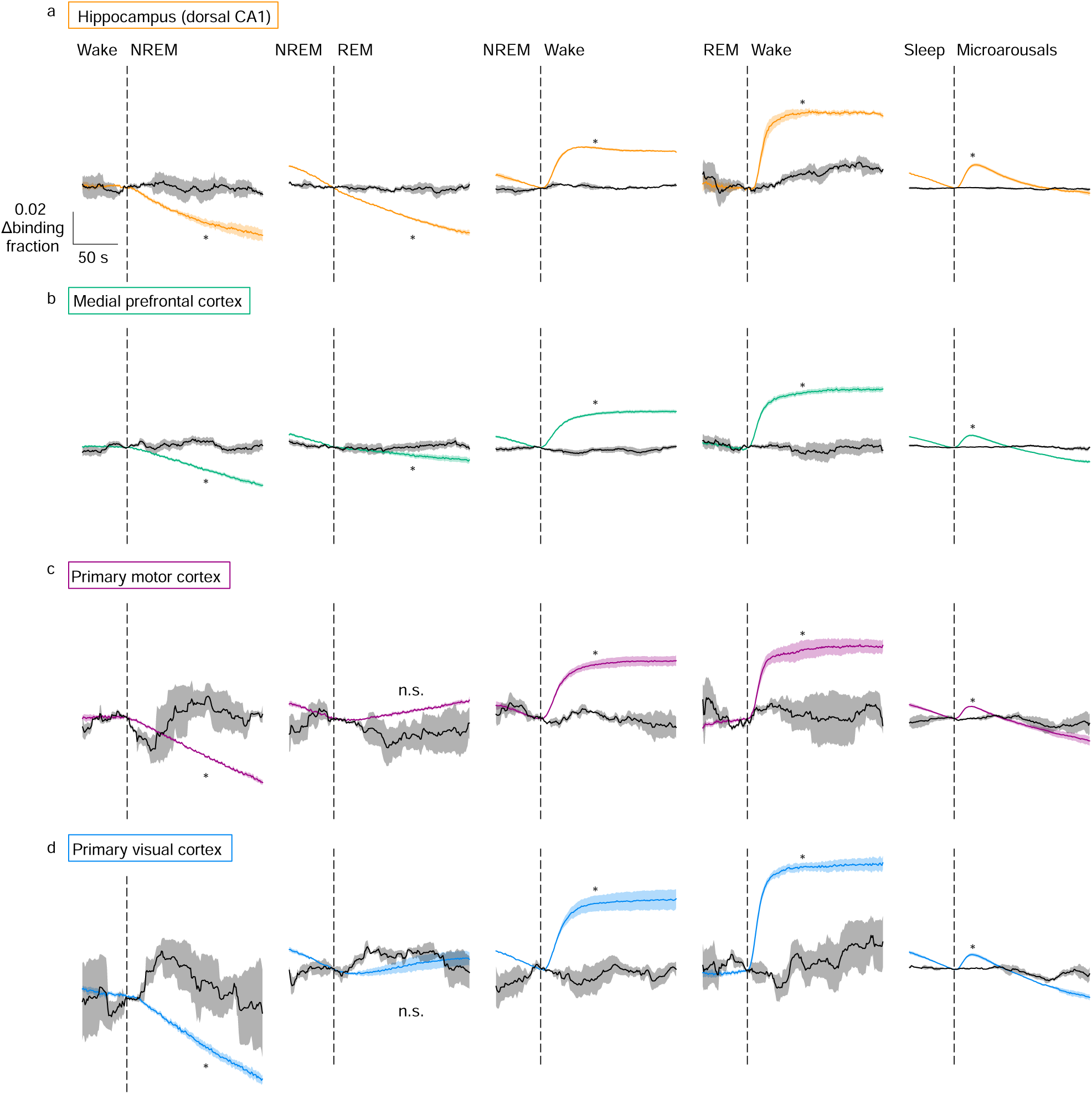
Membrane PKA substrate phosphorylation dynamics are similar across sleep-wake stages in multiple hippocampal-cortical regions. a-d, Behavior state transition-triggered averages of FLIM-AKARm binding fraction (colored traces) and shuffled data (black traces) from (a) hippocampal CA1 (n=3 mice), (b) mPFC (n=18 mice), (c) M1 (n=3 mice), and (d) V1 (n=3 mice). FLIM-AKARm data for (a) are the same as in Fig. 1f, displayed here for comparison. Data show mean ± SEM (paired t-test; FLIM-AKARm vs. shuffled). *: p<0.05; n.s.: not significant, p>0.05. See details in Extended Data Table 1.

The mPFC was selected for further study because it is particularly sensitive to sleep deprivation^67,68^, and excitatory neurons in this region have been implicated in sleep regulation^66^. Moreover, like other brain regions, the mPFC receives neuromodulatory inputs that are differentially released between sleep and wake^42,69^. Given the dense neuromodulatory innervation to this region, we next tested whether these inputs contribute to membrane PKA-SP dynamics. Pharmacological inhibition of noradrenergic and serotonergic signaling, but not cholinergic or adenosinergic signaling, dampened sleep-wake-associated membrane PKA-SP dynamics **(ED Fig. 3)**, indicating that norepinephrine (NE) and serotonin (5-HT) are key upstream regulators.

### Sleep-wake transitions and EEG signatures precede switches in membrane PKA-SP dynamics

Sleep and wake states are characterized by rapid shifts in EEG power across frequency bands. Since membrane PKA-SP dynamics also switch around sleep-wake transitions, we asked whether EEG changes precede or follow the changes in membrane PKA-SP dynamics. First, we calculated mutual information between membrane PKA-SP and EEG power in the delta (0.5-4 Hz), theta (4-8 Hz), sigma (10-15 Hz), and beta (15-30 Hz) frequency bands. We found that changes in EEG power across all frequency bands preceded changes in membrane PKA-SP (**Fig. 3a-d, ED Fig. 4a-d**), suggesting that membrane PKA-SP dynamics are a downstream consequence of sleep-wake state transitions rather than a driver of them, consistent with its role as a reporter of sleep history within each bout.

**Figure 3.**
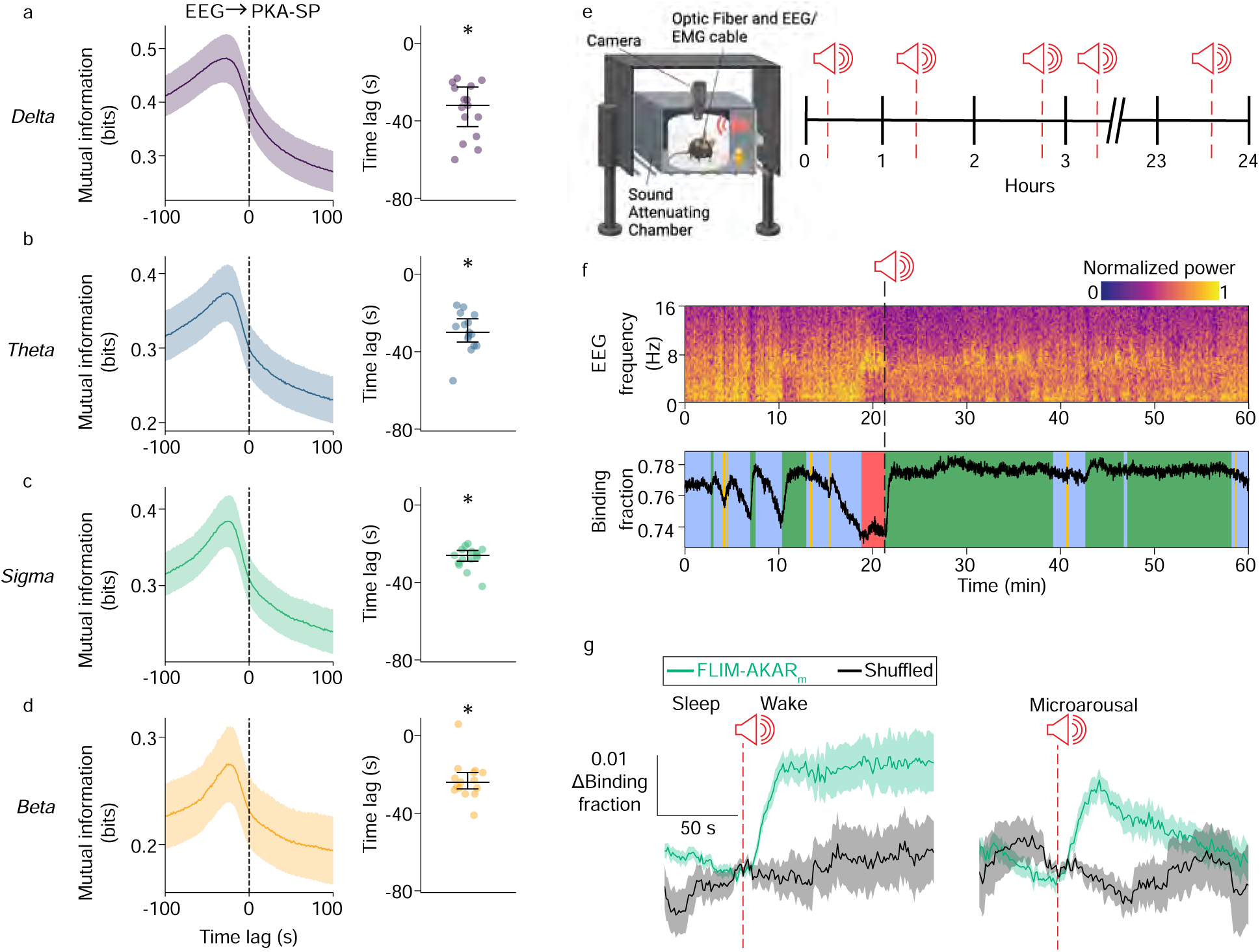
Changes in membrane PKA substrate phosphorylation dynamics follow EEG state shifts and respond to sound-evoked waking. a-d, Average trace (left; mean ± SEM) and peak lag (right; median with interquartile range (IQR)) of mutual information between mPFC EEG band powers and FLIM-AKARm binding fraction. Negative lag indicates EEG power precedes FLIM-AKARm binding fraction (n=15 mice; 1-sample Wilcoxon signed-rank test). e, Schematic illustrating the experiment: during simultaneous EEG/EMG and FLiP recordings, mice were placed in a sound-attenuating chamber. A pure tone was played once per hour at a randomly generated timepoint. f, Example EEG spectrogram and FLIM-AKARm trace with sleep stages. Black dashed line indicates time of tone. g, FLIM-AKARm binding fraction (n=3 mice) and shuffled data aligned to tone onset during sleep when tones led to a wake bout (left) or microarousal (right). Data show mean ± SEM (paired t-test; FLIM-AKARm vs. shuffled). *: p<0.05.

Next, we asked whether membrane PKA-SP dynamics switch following artificial waking by playing sound at random time points (**Fig. 3e**). We observed a rapid increase of FLIM-AKAR_m_ binding fraction following sound-induced waking (**Fig. 3f, g**). Importantly, there was no change in FLIM-AKAR_m_ binding fractions when sound failed to induce waking or was delivered during wake (**ED Fig. 4e**), indicating that membrane PKA-SP responds to the sleep-to-wake transition itself rather than to auditory input. Together, the mutual information analysis and the sound experiment indicate that changes in membrane PKA-SP follow rather than precede EEG power changes and sleep-to-wake state transitions, consistent with membrane PKA-SP encoding sleep history within a bout rather than driving rapid state transitions.

### Membrane PKA-SP dynamics follow asymmetric exponential kinetics

Given the consistent within-bout dynamics of membrane PKA-SP, we asked whether they could be described by a simple mathematical model^50,70^ with parameters that are fixed across bouts. Inspection of the joint distribution of membrane PKA-SP binding fraction and its temporal derivative revealed three distinct dynamical regimes (**ED Fig. 4f**), motivating a mathematical model with three states: a falling phase, a rising phase, and a steady state characterized by high binding fraction and near-zero rate of change. The steady state was identified with a clustering algorithm applied to the joint space (see Methods), and timepoints assigned to this state were treated separately to prevent spurious switching to the exponential models described below. The rising and falling phases were described by two exponential equations with two parameters for each equation (**Fig. 4a**), with the exponential equations taking the same form.

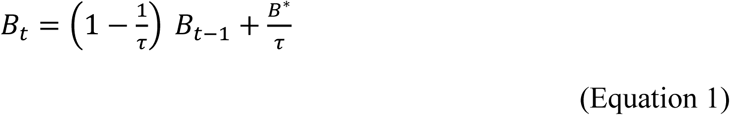

where B refers to FLIM-AKAR_m_ binding fraction at a given time, which exponentially approaches a fixed level (B*) with a time constant τ. During the falling phase (period of PKA-SP decrease), FLIM-AKAR_m_ binding fraction dropped slowly (τ: median = 211.8 s; interquartile range (IQR) = 168.7–234.07 s; n = 21 experiments, 15 mice) to a low asymptote. During the rising phase (period of PKA-SP increase), FLIM-AKAR_m_ increased rapidly (τ: median = 19.1s; IQR = 11.7–23.5 s; n = 21 experiments, 15 mice) and approached a high asymptote. Importantly, parameters optimized on only two to four hours of data, when held fixed, accurately described the full 24-48 hour recording from each animal (**Fig. 4b**, R^2^: median = 0.97; IQR = 0.95-0.98), suggesting that PKA-SP dynamics follow a stereotyped, predictable temporal pattern. Additionally, this exponential model outperformed a linear model for both the rising and falling phases of PKA-SP, as evaluated by Bayesian information criteria (**Fig. 4c-e; see Methods**). Together, these results show that membrane PKA-SP dynamics follow asymmetric exponential kinetics with stereotyped parameters that are consistent across sleep bouts. This exponential form inherently encodes a rate-level relationship with faster dissipation when PKA-SP levels are higher. Crucially, the consistency of the parameters across bouts enables PKA-SP dynamics to provide a reliable and predictable molecular algorithm for tracking proximal sleep history within each bout.

**Figure 4.**
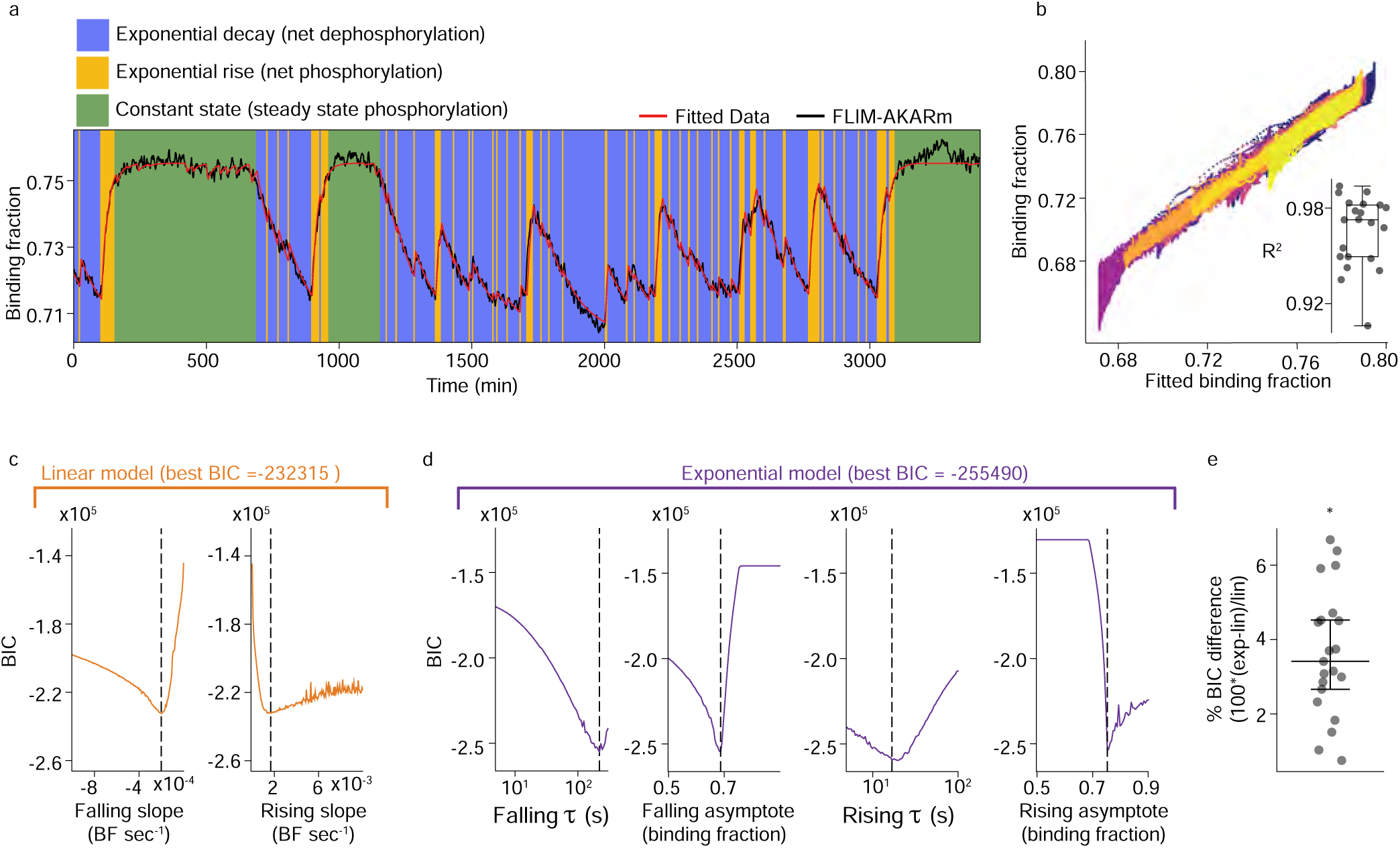
Membrane PKA substrate phosphorylation is accurately modeled with exponential equations. a, Example trace of experimental and fitted data with identified model state. b Scatter plot of experimental vs. fitted FLIM-AKARm binding fraction values across all animals (n=15 mice). Different colors represent data from different mice. Box plot represents distribution of R2 values for all animals (median with IQR; whiskers represent 1.5xIQR). c-d, BIC profile for (c) linear fit model and (d) exponential fit model for an example experiment. Dashed lines represent parameter values for the best overall model based on BIC. e, Percent decrease in BIC from linear to exponential models (n=21 experiments from 15 mice; 1-sample Wilcoxon signed-rank). *: p<0.05. Data show median and IQR.

### Membrane PKA-SP tracks the likelihood of waking

The dissipation dynamics of membrane PKA-SP during sleep suggests that it acts as a molecular timer of proximal sleep history on a bout-to-bout level. Within each sleep bout, we observed that regardless of NREM sleep duration, PKA-SP dissipated gradually (**Fig. 5a**). In contrast, SWA, a brain signal associated with sleep need^5,6,70,71^, did not exhibit significant dissipation during the course of a NREM bout (**Fig. 5b**). This suggests that PKA-SP is a more reliable reporter of proximal sleep history within a sleep bout.

**Figure 5.**
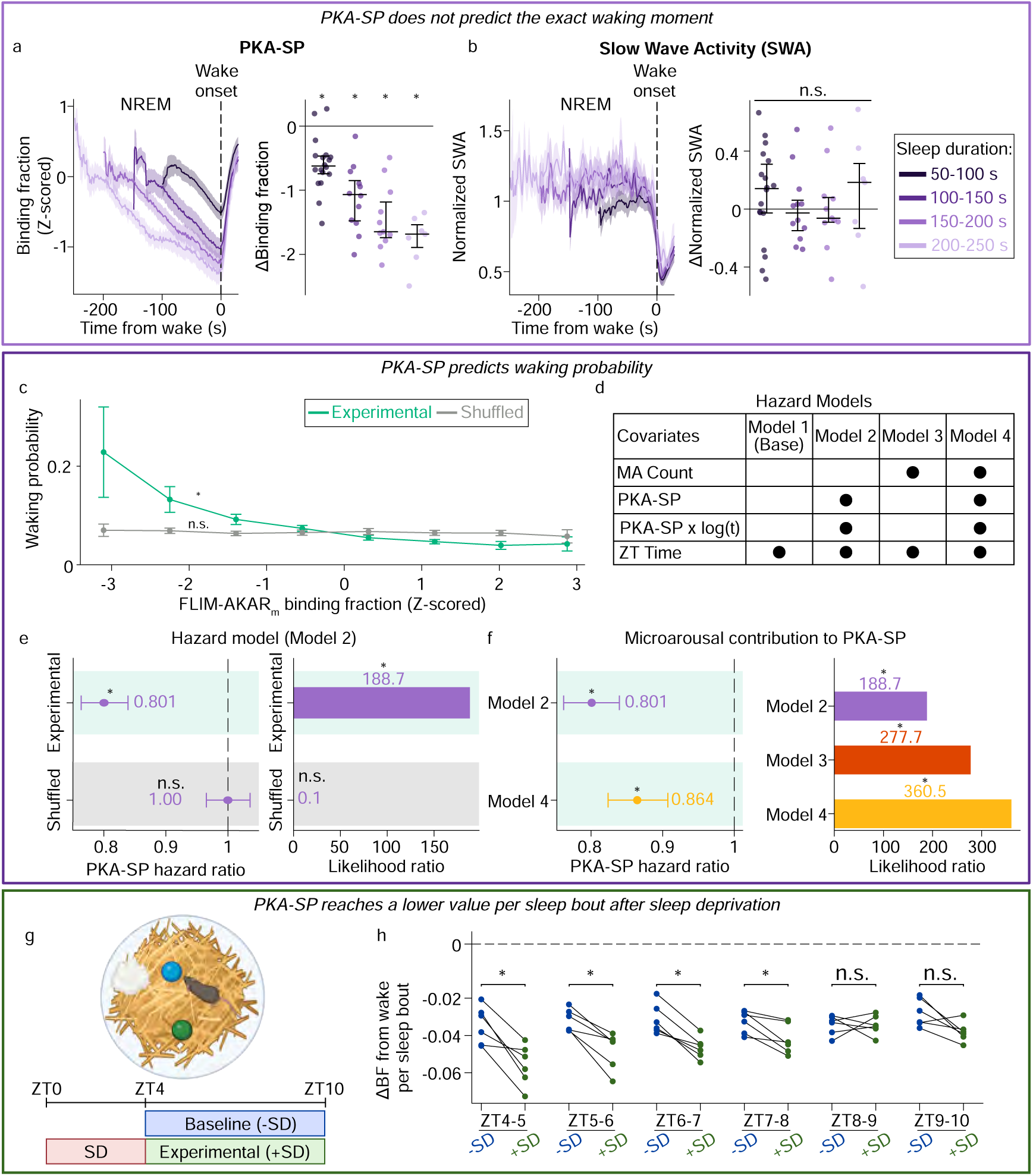
Membrane PKA substrate phosphorylation dynamics track probability of waking. a, Average trace (left; mean ± SEM) and summary of change (right; median with IQR) of FLIM-AKARm binding fraction per bout during NREM aligned to wake onset, binned by the duration of NREM bouts (n=18 mice; 1-sample Wilcoxon signed-rank per bin) b, Average trace (left; mean ± SEM) and summary of change (right; median with IQR) of SWA during NREM aligned to wake onset, binned by the duration of NREM bouts (n=18 animals; 1-sample Wilcoxon signed-rank per bin). c, Waking probability across Z-scored FLIM-AKARm binding fraction in the mPFC during sleep (including NREM, REM, and microarousals). Data show mean ± SEM (n=15 mice). A linear mixed-effects model was fitted to log (waking probability) with binding fraction bin as a predictor; the slope coefficient was tested against zero using a Wald test (see Methods). d, Table summarizing covariates across hazard models (see Methods for details). e, Cox proportional hazard models assessing whether FLIM-AKARm binding fraction predicts instanta-neous waking probability. (Left) Hazard ratios (HR) for FLIM-AKARm binding fraction (Model 2) and shuffled controls (n=18 mice; shown with 95% confidence intervals; Wald test against null HR=1). Binding fraction HR was significantly different from 1; shuffling abolished the effect. Dashed line indicates HR=1 (no effect). (Right) Likelihood ratio comparing Model 2 to the base model (Model 1) (n=18 mice; likelihood ratio test; see Methods). f, (Left) HR for FLIM-AKARm binding fraction from Models 2 and 4 (n=18 mice; shown with 95% confidence intervals; Wald test against null HR=1). (Right) Likelihood ratios for Models 2-4 versus the base model (Model 1; n=18 mice; likelihood ratio test; see Methods). Model 2 reproduced from Fig. 5d for comparison. g, Schematic of procedure for sleep deprivation (SD) by novel object exposure. h, Per-animal mean change in FLIM-AKARm binding fraction in mPFC between the average wake level (ZT4-ZT10) and the end of each sleep bout (including NREM, REM, and microarousals), comparing animals with and without SD (n=6 mice; −SD vs. +SD; Wilcoxon signed-rank test on per-animal means). *: p<0.05; n.s.: not significant, p>0.05.

Because membrane PKA-SP encodes continuous information about proximal sleep history and sleep-wake transitions are probabilistic, we hypothesized that membrane PKA-SP would track the moment-to-moment waking propensity at the bout level. Accordingly, we predicted that PKA-SP would not specify the exact timing of sleep-wake transition, but instead would be associated with waking probability. Indeed, waking occurred at markedly different PKA-SP levels, indicating that PKA-SP during sleep did not predict the precise timing of individual sleep-to-wake transitions (**Fig. 5a**). To test whether PKA-SP tracks the likelihood of waking, we investigated how the probability of waking varied with FLIM-AKAR_m_ binding fraction values during continuous sleep bouts consisting of NREM, REM, and microarousals. Remarkably, the probability of waking decreased gradually and continuously with increasing binding fraction values (**Fig. 5c**). This pattern did not hold with either shuffled data (**Fig. 5c**) or in the presence of PKI (**ED Fig. 5a**), indicating the specificity of the relationship to membrane PKA-SP.

The correlation between membrane PKA-SP and waking propensity could represent the predictive power of PKA-SP itself, or it could simply reflect its co-variation with sleep duration which independently influences waking. To test whether membrane PKA-SP predicts instantaneous waking probability within bouts beyond what is predicted by sleep duration, we built a hazard model with three time-varying covariates **(Model 2, Fig. 5d)**: FLIM-AKAR_m_ binding fraction (z-scored within each recording session), a binding fraction × log(time-in-bout) interaction term to accommodate the time-varying hazard ratio of binding fraction^72^, and Zeitgeber time (ZT) to account for circadian regulation. The hazard ratio for z-scored FLIM-AKAR_m_ binding fraction was 0.801 (significantly different from 1; **Fig. 5e**), indicating that higher binding fraction is associated with reduced waking probability. Specifically, at a fixed sleep duration and circadian time, each standard deviation increase in FLIM-AKAR_m_ binding fraction during sleep corresponded to a 19.9% decrease in instantaneous waking probability. Further, FLIM-AKAR_m_ binding fraction showed a high likelihood ratio, which indicates that it substantially improved the performance of the hazard model. These model effects were abolished with shuffled data **(Fig. 5e)**. Importantly, excluding Zeitgeber time from the model did not alter the hazard ratio for FLIM-AKAR_m_ binding fraction **(ED Fig. 5d)**, indicating that the predictive effect of PKA-SP on instantaneous waking probability is statistically separable from circadian time. Further, to complement the hazard model, a logistic regression model predicting imminent waking showed that FLIM-AKAR_m_ binding fraction held more explanatory power than sleep duration (**ED Fig. 5b**), and this effect was robust across a range of time windows used to define imminent waking **(ED Fig. 5c)**.

We subsequently investigated what membrane PKA-SP could be encoding to facilitate its predictive power of waking probability. We hypothesized that this could be explained by the dissociation between FLIM-AKAR_m_ binding fraction and sleep duration during microarousals. During these brief sleep interruptions within a continuous bout of sleep, FLIM-AKAR_m_ binding fraction rapidly rises while sleep duration continues to increase (**Fig. 1e, f**). Thus, two time points with identical sleep durations can have markedly different FLIM-AKAR_m_ binding fraction levels depending on the occurrence of microarousals. Consequently, PKA-SP can predict waking probability better than sleep duration alone. To test this hypothesis, we built another hazard model **(Model 4, Fig. 5d)** with cumulative microarousal count within a sleep bout as an additional time-varying covariate. This shifted the FLIM-AKAR_m_ binding fraction hazard ratio toward the null (from 0.801 to 0.864; **Fig. 5f**), corresponding to a 34% attenuation in the log-hazard ratio **(ED Fig. 5e)**. This indicates that microarousal history partially accounts for the predictive power of PKA-SP for waking probability. This was further supported by the likelihood ratio tests: either FLIM-AKAR_m_ binding fraction (Model 2) or microarousal history (Model 3) improved the hazard model beyond sleep duration; together (Model 4), they improved the model even further **(Fig. 5d, f)**. Importantly, the contribution of microarousals to the predictive power of PKA-SP held true for a wide range of definitions for duration threshold for microarousals, from 12 to 44s, but became negligible when the microarousal threshold exceeded this range **(ED Fig. 5e, f)**. Together, these results indicate that PKA-SP dynamically integrates sleep duration and interruption to encode bout-level waking probability.

### Membrane PKA-SP reaches lower levels at the end of sleep bouts following sleep deprivation

Thus far, our data show that during natural sleep-wake behavior, membrane PKA-SP dynamics integrate sleep duration and microarousal history to track moment-to-moment waking probability within sleep bouts. To further test whether PKA-SP dynamics are sensitive to sleep history, we asked how membrane PKA-SP responds under conditions of elevated sleep need following sleep deprivation (SD). We performed four hours of SD by novel object introduction (ZT0-ZT4; **Fig. 5g**), a gentle, largely stress-free method of SD^73^. We found that FLIM-AKAR_m_ binding fraction in the mPFC did not significantly increase during the SD period (**ED Fig. 5h),** consistent with the relatively stable PKA-SP levels observed during wake (**Fig. 1e, f**). In contrast, FLIM-AKAR_m_ binding fraction declined significantly more from wake levels within individual sleep bouts following SD than during circadian-matched periods without SD (**Fig. 5h**). This finding was consistent regardless of whether microarousals were classified as part of the sleep bout (**Fig. 5h**) or as distinct wake events (**ED Fig. 5g**). Thus, following SD, PKA-SP dissipated to a lower level after each sleep bout compared to baseline, even though absolute PKA-SP levels during wakefulness did not increase across the SD period. Given that dissipation magnitude increased following SD, when sleep need is presumed to be higher, these results suggest that membrane PKA-SP does not encode the absolute levels of sleep need, but rather tracks how much sleep need has been relieved within each sleep bout, with greater dissipation when more sleep need has accumulated.

## Discussion

Our results show that membrane PKA-SP integrates sleep duration and microarousal history to track moment-to-moment waking probability (**ED Fig. 6**). Fulfilling the criteria established earlier for a molecular signal that tracks within-bout sleep history, membrane PKA-SP: (1) dissipates exponentially throughout each sleep bout with consistent parameters (**Fig. 1, 4**); (2) integrates sleep duration and microarousal history (**Fig. 5**); (3) predicts subsequent waking probability better than sleep duration alone (**Fig. 5, ED Fig. 5**); and (4) shows greater dissipation per bout following sleep deprivation (**Fig. 5, ED Fig. 5**), when accumulated sleep need is higher. Our data additionally reveal features consistent with PKA-SP serving as a molecular integrator of sleep history: its dynamics are conserved across many cortical and hippocampal regions (**Fig. 2**) and are regulated by neuromodulators that are differentially released between wake and sleep and implicated in sleep regulation^1,31,32,42,44,45,69^ (**ED Fig. 3**). Thus, we report the first molecular signal whose in vivo dynamics encode proximal sleep history at the within-bout level.

Membrane PKA-SP predicts moment-to-moment waking probability throughout a sleep bout, providing a molecular correlate of within-bout arousal dynamics at a timescale not captured by existing signals. By dissipating gradually during sleep and rising transiently in response to microarousals, membrane PKA-SP integrates both sleep duration and microarousal history through the biophysical properties of a single molecular signal, predicting waking probability more faithfully than sleep duration alone (**Fig. 5**). Importantly, PKA-SP does not deterministically predict the precise timing of individual wake transitions but rather continuously tracks the probability of waking throughout a sleep bout (**Fig. 5**). While prior work has successfully identified cell types and circuits that promote sleep-to-wake transitions^1,2,20–25^, what continuously encodes the likelihood of waking at any given moment during sleep has remained elusive. PKA-SP provides a molecular signal that does not specify when transitions occur (**Fig. 5a**), but continuously encodes how likely they are throughout each bout (**Fig. 5c-f**). We predict that PKA-SP sets the permissive landscape for waking by gradually increasing waking probability as it dissipates, while superimposed infraslow oscillations in arousal signals, such as NE and 5-HT^32,42,44,74,75^, interact with it to determine the precise moment of transition, consistent with PKA-SP as a downstream signal of neuromodulatory inputs (**ED Fig. 3**).

PKA-SP dynamics are consistent with multiple functional roles. The transient rise in both membrane and intracellular PKA-SP during microarousals is consistent with the dynamics of cAMP during sleep and may support synaptic plasticity and memory consolidation^60,61,64,76^. The gradual within-bout decrease is consistent with homeostatic downscaling^35,59,77^. Most compellingly, integrating the waking probability analysis (**Fig. 5**) with the gradual dissipation dynamics, our data suggest that membrane PKA-SP tracks the relative relief of sleep need within each bout, operationalized here as waking probability since sleep need cannot be measured directly. By exhibiting gradual dynamics throughout sleep, the key kinetic criterion for a molecular substrate of sleep need dissipation, membrane PKA-SP provides the first direct in vivo evidence for a molecular substrate whose existence has been theorized for decades^4,51,70,78^. Beyond this primary kinetic criterion, PKA-SP also fulfills supporting predictions: its dynamics respond to sleep deprivation when sleep need is increased (**Fig. 5**), and its dynamics are modulated by NE and 5-HT (**ED Fig. 3**), two neuromodulators implicated in sleep need regulation^45,79^.

Our results diverge from the classical sleep need signal in two key ways. First, membrane PKA-SP encodes the amount of sleep need dissipated within a bout rather than the absolute level of remaining sleep need. PKA-SP gradually dissipates during sleep but quickly reaches a high plateau during wake (**Fig. 1**), suggesting it resets at wake onset instead of tracking progressive sleep need accumulation^56,80^. Consistently, sleep deprivation increases the drop in PKA-SP at the end of a bout relative to wake without altering absolute PKA-SP level during wake (**Fig. 5, ED Fig. 5**), suggesting that membrane PKA-SP encodes relative relief of sleep need rather than its absolute magnitude. Second, whereas classical models conceptualized sleep need dissipation across bouts over hours^4^, with SWA as the best available marker^5,6^, PKA-SP reveals within-bout organization, a timescale that is functionally important yet has only recently become accessible at the molecular level. SWA remains largely flat within individual bouts^30^, whereas PKA-SP dissipates continuously (**Fig. 5**). The consistent dissipation parameters across bouts (**Fig. 4**) suggest a stereotyped computation of relative relief. We propose that SWA and PKA-SP track sleep need dissipation over long and short timescales, respectively, with bout-level PKA-SP dynamics potentially feeding into the longer-timescale signals that track cumulative sleep need across bouts and days.

Our data provide the most direct evidence to date for an in vivo molecular signal whose dynamics are consistent with bout-level sleep need dissipation. However, whether membrane PKA-SP dynamics drive sleep need dissipation, homeostatic plasticity^35,59,81^, and/or Hebbian plasticity^60^ remains to be established. Although prior data manipulating PKA and its cognate phosphatases^50,54,82^ are consistent with the hypothesis that PKA-SP dynamics regulate sleep need dissipation, the most rigorous test requires closed-loop, real-time perturbation of PKA-SP dynamics during sleep. We are currently developing light-activated PKA inhibitors^83^ to enable these experiments in the future. Importantly, a molecular signal that reliably predicts waking probability has significant value for understanding sleep structure and developing therapeutic targets, independent of its causal role in sleep need dissipation.

Beyond the field of sleep, these results provide a foundation to understand how biochemical properties give rise to emergent properties important for behavior^84^. To our knowledge, the gradual within-bout dissipation dynamics of membrane PKA-SP are not shared by any other characterized signal^31,32,42,44,64,69,69,74,85^. Even PKA-SP itself does not share these dynamics in the intracellular compartment (**Fig. 1**), suggesting that membrane and intracellular PKA-SP may serve distinct functions during sleep. These distinct dynamics likely reflect differential enrichment of PKA, phosphatases, anchoring proteins, or neuromodulator receptors in each compartment, enabling the same molecular signal to encode different information depending on its subcellular location^49^. More broadly, these findings suggest that subcellular compartmentalization may be a general strategy by which neurons efficiently encode distinct behavioral variables using the same molecular machinery.

In conclusion, membrane PKA-SP dynamics reveal a molecular mechanism for within-bout integration of sleep duration and interruption, continuously tracking the moment-to-moment probability of waking (**ED Fig. 6)**. These findings provide the first molecular anchor for sleep need dissipation at the bout level, demonstrate how biochemical kinetics can encode integrated behavioral history on a slow time scale, and identify a potential therapeutic target for sleep disorders such as insomnia, where sleep bout duration and waking probability are dysregulated.

## Methods

### Animals

All aspects of mouse husbandry and surgery were performed following protocols approved by Washington University Institutional Animal Care and Use Committee and in accordance with National Institutes of Health guidelines. The experiments were performed according to the ARRIVE guidelines. All mice in this study were *B6.Cg-Tg(Camk2a-cre)T29-1Stl/J* (Jax: 005359)^62^. All mice were aged between 3 and 10 months at the time of recording. Datasets were composed of both male (n=26) and female (n=16) mice. After surgery, female mice were group-housed with littermates and male mice were singly housed to prevent aggression. All mice were kept in a 12-hour light and 12-hour dark cycle.

### DNA plasmids

The constructs AAV-FLEX-FLIM-AKAR, AAV-FLEX-FLIM-AKAR^T391A^ and AAV-FLEX-PKIα-IRES-mRuby2 were described previously^8^ (Addgene #s 60445, 60466 and 63059). AAV-DIO-EF1α-mKate-biPAC^65^ is a gift from Drs. Shiqiang Gao and Georg Nagel at University of Wuerzburg, Germany. AAV-FLEX-FLIM-AKAR_m_ was constructed by the addition of a farnesylation domain to the C-terminus of AAV-FLEX-FLIM-AKAR (Genscript), and will be deposited to Addgene.

### Viral production and mouse surgery

AAV2/1-FLEX-FLIM-AKAR-SV40 was packaged by the Boston Children’s Hospital Viral Vector Core and used at 1.05×10^13^ gc/mL. AAV2/1-DIO-EF1α-mKate-biPAC was packaged by Neurotools and used at 1.84×10^12^ gc/mL. AAV2/1-FLEX-FLIM-AKAR_m_, AAV2/1-FLEX-AKAR^T391A^, and AAV2/1-FLEX-PKIα-IRES-nls-mRuby2 were packaged by the Washington University Viral Vector Core and used at 1.4×10^12^ gc/mL, 2.05×10^12^ gc/mL, and 2.16×10^12^ gc/mL, respectively.

For AAV viral injection, we targeted multiple brain regions with the following stereotaxic coordinates relative to Bregma. Dorsal CA1: anteroposterior (AP) −1.78 mm; mediolateral (ML) −1.58 mm; dorsoventral (DV) 1.36 mm ventral to the pial surface 2) medial prefrontal cortex (mPFC): AP +1.90 mm; ML −0.15 mm; DV 2.0 mm 3) motor cortex (M1): AP +1.85 mm; ML −1.5 mm; DV 0.25 mm 4) visual cortex (V1): AP −3.0 mm; ML −2.0 mm; DV 0.3 mm. For all injections, 500 nL of total virus was delivered at a rate of 50 nL/min using a UMP3 micro-syringe pump (World Precision Instruments) with a glass pipette. After viral injection, a fiber-optic cannula (Doric Lenses, MFC_200/245-0.37_2.5mm_MF1.25_FLT) was implanted 0.1 mm dorsal to viral injection site as described previously^63,86^. EEG and EMG implants (Pinnacle, 8402-SS) were placed as previously described^63^.

### Fluorescence Lifetime Photometry (FLiP), EEG, EMG recordings, and analysis

A FLiP setup was built as previously described^63,86^. FLiP (1 Hz), EEG (400 Hz), EMG (400 Hz), and video recordings (25 frames per second) were performed as described previously^63^ except for the following: each recording was allowed to run continuously for 6-72 hours. Behavior was monitored with synchronized video recordings through Bonsai (https://bonsai-rx.org/) which were processed post-hoc using DeepLabCut (DLC) for movement analysis. Calculation of

FLIM-AKAR binding fraction was performed using a custom software package as described previously^86^. Binding fraction is *P_FRET_* from the fitted double exponential curve.

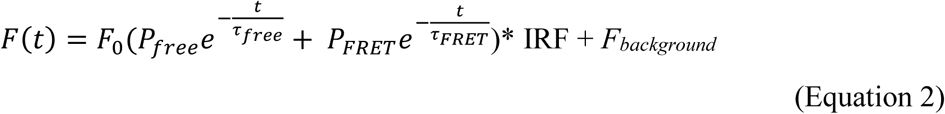

where F(t) is the photon count at photon arrival time t, F_0_ is the peak photon count, τ_free_ and τ_FRET_ are the fluorescence lifetime decay constants of donors that are free and donors that have undergone FRET, respectively, and P_free_ and P_FRET_ are the corresponding fractions. τ values are 2.14 ns (free) and 0.69 ns (FRET) for FLIM-AKAR_m_^8^. For determining sleep stages, EEG and EMG data were downsampled to 200 Hz before sleep-wake behavior was scored using a custom Python package as described previously^63^.

### Histology

The location of viral expression and fiber implant was assessed with histology after recordings. Tissue fixation and slicing were performed as previously described^86,87^ except for the following. Sections were stained with Hoechst before being mounted on glass slides (VWR cat. no. 48311-702) using Fluoromount-G^TM^ mounting media (Invitrogen Cat. no. 00-4958-02). Sections were imaged using an epifluorescence microscope (Nikon E800) with a camera (Teledyne Photometrics, CoolSnap EZ) and the software QCapture Pro.

### Confocal microscopy

Fixed hippocampal brain slices were imaged on a Nikon Ti2 microscope with a CSU-W1 SoRa spinning disk confocal head equipped with a Hamamatsu Fusion BT sCMOS camera through the Washington University Center for Cellular Imaging (WUCCI). A 20x, 0.75 NA air objective (axial optical resolution 1.5 µm) (Nikon) was used for capturing subcellular expression patterns in the hippocampus. These images are displayed as a single optical section **(Fig. 1b; left and middle)**. For higher resolution imaging of dendritic spines, a 60x, 1.40 NA oil immersion objective and 4x SoRa relay lens were used (axial optical resolution 0.5 µm). Z-stacks were acquired at 0.2 µm steps and displayed as a maximum intensity projection across 15 slices (**Fig. 1b; right**). Fluorescence was excited at 488 nm (FLIM-AKAR and FLIM-AKAR_m_) and 405 nm (Hoechst stain) laser lines and collected through 535/40 nm and 460/50 nm bandpass emission filters, respectively.

### Sound-triggered waking

To determine whether waking causally increases PKA-SP, the recording chamber was placed in a sound-attenuating box to limit external sounds. A custom MATLAB software was used to generate a 5-second tone at a frequency of 8 KHz at a random time each hour. Continuous microphone recordings were collected through Bonsai (https://bonsai-rx.org/) to record sound and align the time of tone delivery to the photometry data.

### In vivo pharmacology

To understand the role of neuromodulator signaling in mediating PKA-SP, we administered propranolol (20 mg/kg), prazosin (1 mg/kg), scopolamine (5 mg/kg), DPCPX (2 mg/kg), MDL100907 (5 mg/kg), or vehicle (either saline or DMSO in saline) during simultaneous FLiP and EEG/EMG recording. Drug was delivered via intraperitoneal (i.p.) injection every 2 hours throughout a 6-12 hour window during the light period (ZT0-ZT6, ZT6-ZT12, or ZT0-ZT12). All analysis comparing drug and vehicle conditions were matched for circadian time.

### Sleep deprivation (SD) and data analysis

To understand the effect of SD on PKA-SP dynamics, mice were kept awake for 4 continuous hours (ZT0-ZT4) during simultaneous FLiP and EEG/EMG recordings. SD was enforced using novel object introduction^73^. Objects were placed and removed with minimal disruption of the mouse, the cage, and the nest. Prior to the deprivation experiments, objects were rinsed with water, dish soap, and KennelSol and then dried before use. Mice were acclimated to the experimental cage for a minimum of 12 hours before baseline recording and then recorded during ZT0-ZT10 on the day before SD and the day of SD.

SD data were compared against circadian-matched control data recorded the day before SD. To quantitate the change in binding fraction across sleep bouts (including NREM and REM), we computed the difference between the binding fraction at the end of each bout and the mean binding fraction across all wake epochs during ZT4-ZT10 (i.e. the period after SD). Microarousals within sleep bouts were either treated as sleep (**Fig. 5**) or as wake (**ED Fig. 5g**). Change in binding fraction was averaged across bouts within each animal, and a Wilcoxon signed-rank test was applied to the resulting per-animal means to evaluate whether post-SD bouts differed from circadian-matched controls.

### biPAC stimulation

In order to test whether FLIM-AKAR_m_ binding fraction reached a plateau during wake due to ceiling effects of the sensor, we co-expressed Cre-dependent light activated adenylate cyclase (biPAC)^65^ and Cre-dependent FLIM-AKAR_m_ in excitatory neurons in the mPFC of *Tg(Camk2a-cre)* mice. During a waking bout, after FLIM-AKAR_m_ binding fraction reached a plateau, we stimulated biPAC using a pulse of 470 nm light with a power of 100.5μW for variable durations between 10 ms and 1000 ms.

### Mathematical modeling of PKA-SP dynamics

To characterize the temporal dynamics of the signal, we modeled the time series using a three-state dynamical framework. FLIM-AKAR_m_ binding fraction, B(t), was fit locally in sliding windows by alternating between two exponential equations representing distinct dynamical regimes. Both were described by Equation 1, but the two equations had different *_ττ_* and B*. This formulation is equivalent to a first-order autoregressive (AR(1)) process, or an exponential equation. We also explored an alternative linear model (**Fig. 4**) and compared the accuracy of both models. Both models shared the same sliding-window fitting procedure described below; they differed only in whether the rise and fall within each state were described by exponential or linear functions.

Model fitting was performed in sliding windows with a length of 40 data points and a step size of 39 data points, allowing local dynamics to be inferred while maintaining continuity of the fitted trajectory. We used 2-4 hours of observed data containing >=80% sleep behavior for fitting. These criteria allowed for the data to contain a sufficient number of both falling and rising periods to fit the most accurate parameter values without overfitting. Within each window we evaluated all possible state trajectories containing up to two switches between the two dynamical regimes. This number was chosen to capture the empirically observed slow-fall and fast-rise dynamics of PKA-SP while preventing overfitting to high-frequency noise. For each candidate trajectory, the initial value was set to the final value of the previous time window. The trajectory minimizing the root-mean-square error (RMSE) relative to the observed data was selected as the optimal description of the local dynamics. Note that RMSE serves here purely as a local criterion for identifying the best-matching trajectory from the candidate set defined by the model parameters and is distinct from the global model evaluation described below. The resulting procedure produced a reconstructed trajectory B(t) and a sequence of inferred dynamical states s(t)∈{1,2}, where 1 and 2 represent the falling and rising states, respectively.

To quantify model quality while penalizing excessive switching, we computed a Bayesian Information Criterion (BIC)

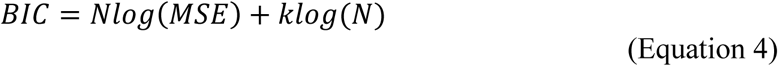

where N is the number of samples in the time series, MSE is the mean squared residual error, and k is the effective number of model parameters, defined as the number of detected state switches plus 4 (the number of parameters describing the autoregressive processes).

The parameters governing the two dynamical regimes (*_τ_*1, *_τ_*2, B1*, B2*) were determined using a multi-stage optimization procedure designed to efficiently explore parameter space while minimizing the BIC score. First, a coarse global search was performed by randomly sampling candidate parameter sets within predefined bounds. The time constants associated with the two states were sampled logarithmically within biologically plausible ranges (decay: 20–300 s; rise: 5–60 s), while the asymptotic values (B1*, B2*) were sampled uniformly within the interval 0.6–0.85 (**Fig. 1e, f**). Each candidate parameter set was evaluated by the BIC score. To accelerate the coarse search, the BIC was initially evaluated using a random subsample of the time series. The best-performing candidate parameter sets from this stage were then re-evaluated on the full dataset. Finally, the top candidates were used as initial seeds for a local numerical optimization using a bound-constrained quasi-Newton algorithm (L-BFGS-B) as implemented by package scipy.optimize.minimize [version 1.16.2], which minimized the BIC score with respect to the four model parameters. Parameters corresponding to relaxation time constants were optimized in logarithmic space to improve numerical stability and ensure positivity for parameters that span orders of magnitude. The parameter set yielding the minimal BIC was selected as the optimal description of the sleep-bout dynamics for each recording.

As FLIM-AKAR_m_ binding fractions are at a plateau during wake with fluctuations that may reflect behavior changes, we applied a two-dimensional Gaussian mixture model clustering algorithm (sklearn.mixture.GaussianMixture [version 1.7.2]) to the joint space of binding fraction and its temporal derivative (**ED Fig. 4f**) to capture inertial dynamics of binding fraction activity not described by the two-state exponential model. The signal was first smoothed using a moving-average filter (window = 40 data points) to suppress high-frequency noise that would otherwise distort derivative estimation and the resulting phase space. A 40 second window was chosen because this was the fastest kinetics of observed FLIM-AKAR in vivo^88^. The derivative was computed using the first temporal difference, and both variables were z-scored (ddof = 1) before clustering. A Gaussian mixture model with three components and full covariance matrices was fit using expectation–maximization with multiple random initializations. The cluster characterized by the highest binding fraction and slope closest to zero was identified as a steady-state regime because this largely captured the plateau during wake. Timepoints assigned to this cluster were treated as an independent third dynamical state and replaced by the fitted binding fraction value at the previous timepoint (t−1), enforcing persistence of the steady-state trajectory for each high binding fraction plateau while preventing spurious switching in the model.

### Logistic regression

To identify the relative contributions of predictors to waking events, we employed binary logistic regression with a fixed-effects approach to control for individual animal differences. The dependent variable was a binary indicator of whether the animal would wake within a certain amount of time (16 seconds for **ED Fig. 5b**, variable window as a robustness check in **ED Fig. 5c**). Predictors were FLIM-AKAR_m_ binding fraction values and sleep duration (including NREM, REM, and microarousals). Both predictors were standardized (z-scored) within each experiment to enable direct comparison of effect sizes.

The model was specified as:

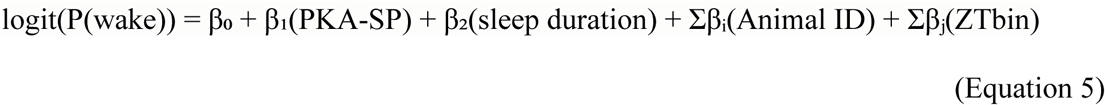

where Animal ID denotes indicator variables for individual animals (with one animal serving as reference) and ZTbin denotes indicator variables for Zeitgeber time grouped into two-hour bins (with one bin serving as reference, 11 indicator variables spanning ZT0-ZT24). This fixed-effects specification controlled for between-animal variability in baseline waking probability and for circadian variation across ZT time, while estimating the associations of PKA-SP and sleep duration with the probability of waking. The model converged successfully under maximum likelihood estimation.

We assessed predictor importance using three complementary approaches: (1) comparison of standardized coefficient magnitudes, (2) Wald tests of individual coefficients with 95% confidence intervals, and (3) likelihood ratio tests comparing full models to reduced models excluding each predictor. Likelihood ratio statistics were calculated as LR = 2(LL_full - LL_reduced), where LL denotes log-likelihood and compared against a chi-square distribution with degrees of freedom equal to the number of parameters removed from the reduced model (df=1 for PKA-SP and sleep duration, each modeled as a single continuous predictor). These analyses were conducted using the package statsmodels [version 0.14.5]. Statistical significance was set at α = 0.05 for all tests.

### Cox proportional hazard model

To assess the relationship between PKA-SP and the instantaneous waking probability during sleep, we employed a Cox proportional hazard model. A counting-process structure was used in which each sleep bout (including NREM, REM, and microarousals) was represented as a series of epochs in start-stop format with timestamps defined as the timepoints at each photometry measurement relative to bout onset. Cumulative sleep duration, and time-varying covariates were updated at each epoch. The event indicator was set to 1 at the final epoch of each bout, the epoch immediately preceding sustained wake (wake lasting longer than microarousal duration threshold) and 0 for all other epochs. If the entire recording ended in a sleep bout, we excluded that bout from analysis due to the uncertainty of the waking time. No censoring was applied as all remaining bouts terminated in a wake event.

Four hazard models in a nested comparison sequence were fit **(Fig. 5d)**, all of which contained a Gaussian shared frailty term on the log-hazard scale to account for between-animal variability in baseline wake propensity as a random effect and a binned Zeitgeber time (ZT) covariate to control for differences in wake propensity based on time of day. ZT bin size was set to 2 hours for this analysis. In robustness checks, varying the bin size between 1 and 4 hours did not substantially alter the hazard ratios for FLIM-AKAR_m_ binding fraction, all of which remained significantly different from 1. Model 1 was a base model containing only ZT bins, serving as the reference for all nested comparisons. Model 2 included ZT bins, FLIM-AKAR_m_ binding fraction, and a FLIM-AKAR_m_ binding fraction × log(time-in-bout) interaction term. Model 3 included ZT bins and a cumulative microarousal count, defined at each epoch as the number of microarousals that had occurred within the current sleep bout up to that point. Model 4 included ZT bins, FLIM-AKAR_m_ binding fraction, cumulative microarousal count, and the interaction term as time-varying covariates. FLIM-AKAR_m_ binding fraction was z-scored within each recording session (24-48 hours), so that the coefficient reflects within-session fluctuations in PKA-SP on the scale of within-session standard deviations. Dependence of waking probability on time already slept in the bout was captured implicitly by the non-parametric baseline hazard function h_0_(t), without requiring duration as an explicit covariate.

The proportional hazard (PH) assumption was assessed via scaled Schoenfeld residuals^72^. In Model 2 without the interaction term, FLIM-AKAR_m_ binding fraction showed a significant departure from PH (χ²=15.27, df=1, p=9.3×10⁻⁵). Because the effect of FLIM-AKAR_m_ binding fraction on waking probability varies with time-in-bout, models containing FLIM-AKAR_m_ binding fraction as a covariate (2 and 4) also incorporated a FLIM-AKAR_m_ binding fraction × log(time-in-bout) interaction term. Importantly, log(time-in-bout) is centered at its mean log value across all epochs, so that the reported FLIM-AKAR_m_ coefficient represents the effect at this time value. With this addition, the global test statistic and the covariate-level test statistic for FLIM-AKAR_m_ for Model 2 was not significant (Global: χ²=15.87, df=13, p=0.256; PKA-SP: χ²=0.011, df=1, p=0.918). However, the covariate-level test statistic for the FLIM-AKAR_m_ binding fraction × log(time-in-bout) itself remained significant (χ²=10.95, df=1, p=9.30×10^-4^), representing a second-order time-varying effect. Since the PH assumption was met globally and for the primary covariate of interest, this residual non-proportionality was not further modeled.

To characterize the relationship between PKA-SP and waking probability and assess the contribution of microarousals to this signal, the following analyses were performed. First, the PKA-SP hazard ratio (HR) and 95% confidence interval were extracted from models 2 and 4, with statistical significance assessed via the Wald test. The attenuation of the PKA-SP coefficient between these two models, which reflects the proportion of PKA-SP’s effect on waking probability shared with microarousal accumulation, was computed as the percentage reduction in the log-hazard ratio.

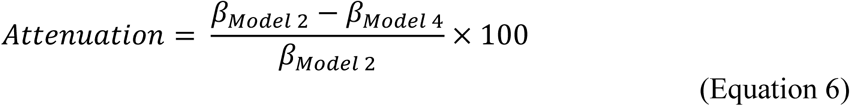

where *_βModel_* _2_ and _β*Model* 4_ are the log-hazard ratios of FLIM-AKAR_m_ binding fraction from models 2 and 4. To test robustness to the definition of the duration threshold of a microarousal, both the HR estimates and the attenuation were computed across all microarousal duration thresholds (12–56 seconds in 4-second increments). Second, likelihood ratio tests (LRT) were used to assess the independent contributions of PKA-SP and microarousal count beyond the base model. Models 2, 3, and 4 were each compared against the base model (Model 1), yielding LRT statistics evaluated against chi-square distributions with 2, 1, and 3 degrees of freedom respectively. This comparison was also repeated across all microarousal duration thresholds. Third, to assess the contribution of circadian time to the PKA-SP signal, the HR of FLIM-AKAR_m_ binding fraction from Model 2 was compared to that from the same model fit without the ZT covariate.

To quantify estimation uncertainty around the attenuation of the PKA-SP hazard ratio upon inclusion of microarousal count, we constructed a 95% bootstrap confidence interval on the attenuation values. To assess whether this attenuation exceeded what would be expected from randomly timed microarousals, we performed a permutation test in which the temporal positions of microarousals within a recording session were randomized while preserving their total number and individual durations. Models 2 and 4 were refitted on each permuted dataset and the attenuation was recomputed. 1000 permutations were performed to build a null distribution of attenuation values. This analysis was repeated across a range of microarousal duration thresholds to determine the threshold above which the observed attenuation was no longer significantly greater than chance. For each threshold, a one-tailed p-value was computed as the proportion of permuted attenuations greater than or equal to the observed attenuation.

Hazard models were fit in R [version 4.5.2] using the coxme package [version 2.2-22], called via subprocess in Python.

### Quantification and statistics

Further descriptions of quantifications and statistics can be found in figures, figure legends, results, and ED Table 1. Unless otherwise specified, traces over time were shown as mean ± standard error of the mean (SEM) to preserve temporal waveform structure; scalar summary statistics were reported as median with interquartile range (IQR) and non-parametric statistical tests were used.

#### State transition-triggered binding fraction

To calculate state transition-triggered changes in binding fraction, we subtracted the binding fraction value at each behavioral state transition onset from all values, yielding a baseline-corrected measure of change relative to transition onset **(Fig. 1d, f**; **Fig. 2**; **Fig. 3g; ED Fig. 4e)**. For drug experiments **(ED Fig. 3)**, binding fraction in both the drug and saline conditions were z-scored together (ddof=1) within each animal. Relative or z-scored binding fraction was extracted from a window of 50 seconds before and 120 seconds after each state transition to capture the full post-transition dynamics. Before-transition behavior bouts lasting less than 50 seconds and after-transition behavior bouts lasting less than 120 seconds were excluded. For each transition type, the mean binding fraction across all instances within an animal was computed to obtain the average transition-triggered activity per animal, and then the mean across all animals was computed for the figures.

Transition-triggered changes were described by amplitude of change, as defined by the mean binding fraction value in a post-transition window minus the value at the behavior state transition. Post-transition windows were defined to capture characteristic features of the sensor dynamics for each transition type, based on observations of the average transition-triggered traces. Post-transition analysis windows were defined separately for each compartment-targeted sensor. For FLIM-AKAR_m_ binding fraction, windows were set to 75-100 s for transitions into NREM and REM (to capture PKA-SP after sufficient dissipation), 50-75 s for transitions into wake (to capture steady-state PKA-SP during wake), and 10-30 s for transitions into microarousal (to capture the peak of the transient change). For FLIM-AKAR, windows were set to 40-65 s for transitions into NREM and REM (to capture PKA-SP at the low plateau), 30-40 s for transitions into wake, and 10-30 s for transitions into microarousal (to capture the peak of the transient change).

When a shuffled control was used for comparison, shuffling was performed by first separating the FLiP data into 40 s bins and then randomizing the order of the bins. 40 s was chosen as the bin size because it approximates the shortest duration of observed PKA-SP transients, preserving local dynamics while disrupting the global relationship between PKA-SP and sleep-wake stages. Sleep-wake staging labels were preserved.

To compare transition-triggered PKA-SP activity between experimental and control groups, we employed a two-stage analysis that accounts for the hierarchical structure of the data, in which individual transition instances are nested within animals and animals are nested within groups. In the first stage, a Huber T M-estimator^89^ was used to generate a robust mean per animal per transition type. This approach applies ordinary least-squares weighting to observations within c = 1.345 standard deviations of the estimated mean and linearly downweights observations beyond this threshold, reducing the influence of outlier transition instances. In the second stage, group differences were assessed by applying either a Welch t-test (which does not assume equal variances between groups, used when comparison groups were composed of different animals) or a paired t-test (used when comparison groups contained the same animals) to the per animal robust means. The normality of the distribution in the per animal means was tested and confirmed for the mPFC data; normality was assumed for other brain regions without formal testing. Analyses were conducted using the statsmodels package [version 0.14.5].

#### EEG spectral analysis

EEG power spectral density was estimated using Welch’s method, applying Fast Fourier Transform to 10-second Hann-windowed segments with 90% overlap. Frequency bands were defined as: delta (0.5-4 Hz), theta (4-8 Hz), sigma (10-15 Hz), and beta (15-30 Hz). In **Fig. 5b**, delta power during NREM sleep was normalized to the mean delta power across all NREM epochs. All spectral analyses were performed using the package scipy.signal [version 1.16.2].

#### Mutual Information Analysis

To determine whether changes in EEG frequency band power precede or follow changes in PKA-SP, we computed time-lagged mutual information (MI) between each EEG frequency band and FLIM-AKAR_m_ binding fraction across the full recording (24-48 hours). Raw binding fraction and EEG power values were used directly without normalization. MI values were estimated using a histogram-based approach. Joint probability distributions of FLIM-AKAR_m_ binding fraction and EEG power were estimated using a 100×100 bin grid (10,000 bins total), with bin boundaries for each variable determined independently by the range of observed values within each experiment, resulting in experiment-specific bin widths. Lagged MI was computed at 1-second intervals up to a maximum lag of 250 seconds in both directions to capture the peak and decay of the MI. The peak lag was identified per experiment from the MI-vs-lag curve. A one-sample Wilcoxon signed-rank test was used to assess whether peak lags differed significantly from zero across recordings (α = 0.05).

#### Waking Probability

To assess the relationship between PKA-SP and probability of waking, we z-scored FLIM-AKAR_m_ binding fraction values from within sleep periods only (consisting of NREM, REM, and microarousals) from each recording within each animal. All values across animals were binned into 8 equal-width bins. For each animal we then calculated the proportion of binding fraction values in each bin that occurred within a defined time window before waking (16 seconds in **Fig. 5c**). The mean and standard error of the mean was calculated across animals.

To test whether waking probability exhibited graded exponential change with binding fraction, we fit a linear mixed-effects model with binding fraction bin position as the predictor, and the log-transformed waking probability as the dependent variable. A random intercept for each animal was included to account for between-animal differences in baseline waking probability. The model was specified as:

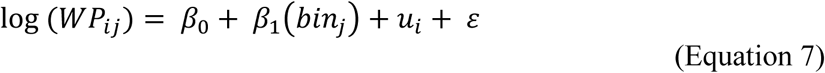

where *WP_ii_* is the wake probability for animal *i* at binding fraction bin position *j*, *β*_0_ is the population-level intercept, *β*_1_ is the fixed effect of binding fraction bin (*bin_j_*), *u_i_* is the random intercept for animal *i*, and *ε* is the residual error. Model parameters were estimated using restricted maximum likelihood (REML). The binding fraction coefficient *β*_1_ was tested against zero using a Wald test (α = 0.05). Models were fit using statsmodels [version 0.14.5].

#### Rate of binding fraction decrease

To quantify the bout-level decay constant of FLIM-AKAR_m_ binding fraction during sleep (**ED Fig. 1e, f)**, we fit an exponential function to each bout of sleep (including NREM and REM) lasting more than 100 s and following a wake bout of greater than 50 s, to ensure the binding fraction starts from a wake-level baseline, comparable to the condition for scopolamine injection

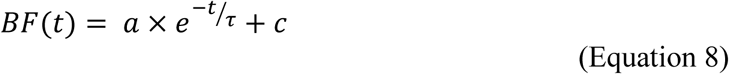

where *BF*(*t*) is binding fraction at time *t*, *τ* is the decay constant, *A* is the initial amplitude above the asymptote, and *c* is the asymptote value. To quantify the decay constant of FLIM-AKARm binding fraction after i.p. injection of scopolamine (5mg/kg) during wake, we applied the exponential fitting in a 100 s window from the time of the injection to capture the full decay dynamics. To account for potential animal-level variability, we collected paired recordings using the same mice for baseline and scopolamine. For both groups of decay constants, bouts were excluded if the data was better fit with a linear function as evaluated by BIC (see above).

Differences in decay constants between conditions were evaluated with a two-stage approach: a Huber T M-estimator was used to compute a robust mean decay constant per animal, and a one-tailed Wilcoxon signed-rank test was applied to the per-animal means, testing whether *τ* was smaller after scopolamine than in controls.

#### Software and Code Availability

All analyses were performed in Python [version 3.11.6] unless otherwise specified. The Python codes for sleep staging are available at https://github.com/YaoChenLabWashU/neuroscience_sleep_scoring. All other analysis code can be found in the following GitHub repository: https://github.com/YaoChenLabWashU/Publication/tree/924f1bbead6b3286c67c722240f2254190131d8f/PKASleep_Manuscript.

#### Data Availability

All data are available upon request.

## Acknowledgments

We thank Paul Taghert, Ed Han, Tim Holy, Wayland Cheng, Gaia Tavoni, Paul Shaw, and all members of Yao Chen’s lab for feedback on this project. We thank Keith Hengen, Daniel Kerschensteiner, Martha Bagnall, Yu Huan, Waleed Babar, Aditi Ashwin, Linda Richards, and Erik Herzog for feedback on this manuscript. We thank Georg Nagel and Shiqiang Gao for sharing the biPAC plasmid. We thank Serena Wang, Carolyn Xiao, and Fapianey Alexandre for help with sleep scoring, and Samarth Aggarwal for adapting DeepLabCut for our video analyses. We thank Peter Bayguinov and Washington University Center for Cellular Imaging (WUCCI) for assistance with high resolution imaging using Nikon Ti2 microscope CSU-W1 SoRa spinning disk confocal. This work was supported by the Hope Center Viral Vectors Core at Washington University School of Medicine.

## Author Contributions

E.T. and Y.C. conceptualized the study. A.F. and W.S. designed and developed the computational model for FLIM-AKAR_m_ binding fraction. E.T. developed the primary computational pipeline including the semi-automated sleep scoring program. E.T., K.G., and

A.O. performed surgery and managed animal care. E.T., K.G., S.M., D.G., and Z.B. performed and analyzed FLiP experiments. K.G. and D.G. designed, performed, and analyzed sleep deprivation experiments. Z.B. designed, performed, and analyzed optogenetic experiments. E.T. and K.G. designed, performed, and analyzed pharmacologic experiments. S.M. designed, performed, and analyzed sound experiments. Y.C. supervised the study. E.T. and Y.C. wrote the manuscript. All authors reviewed the manuscript.

## Supplementary Information is available for this paper

**Extended Data 1.**
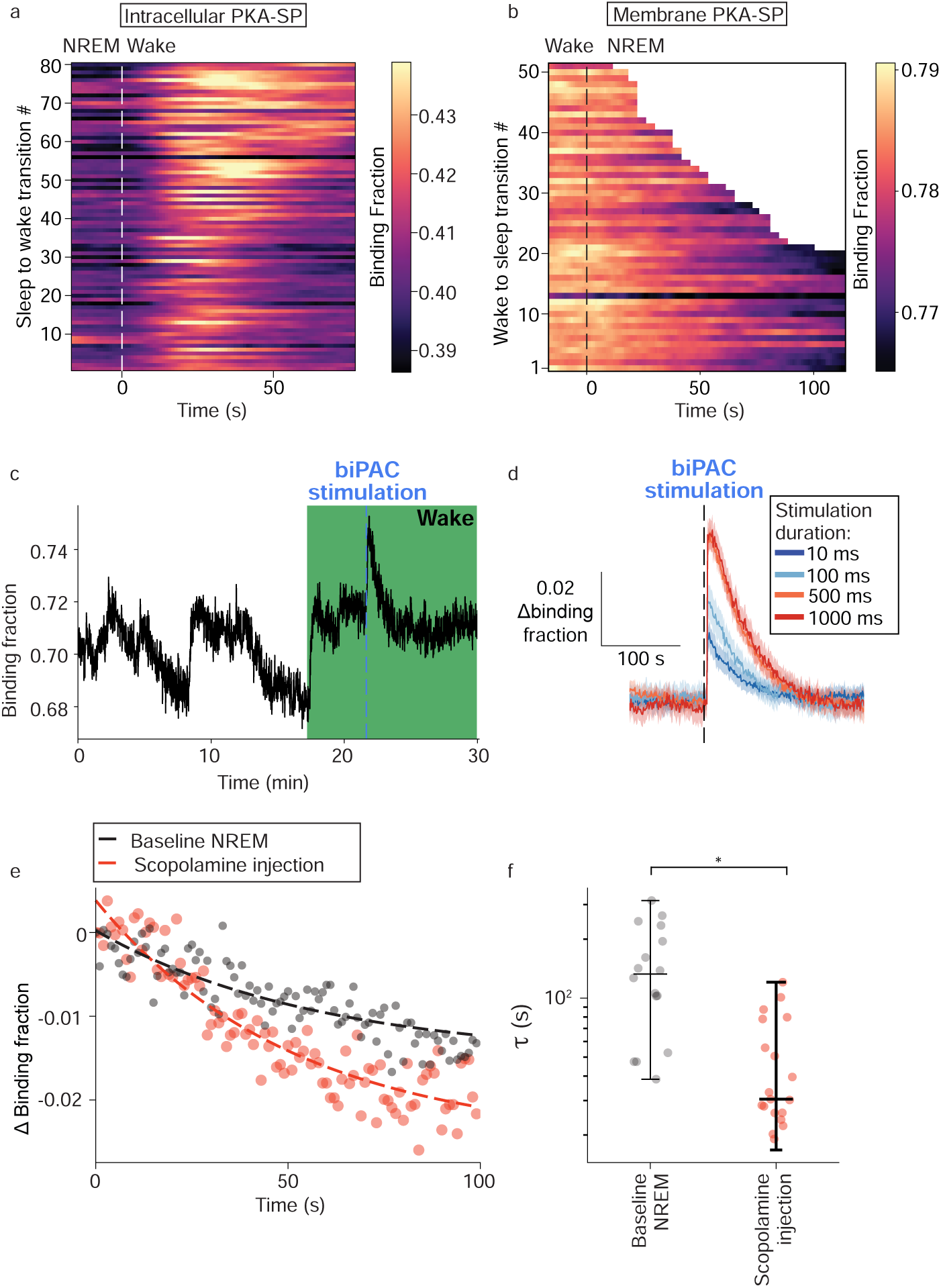
Compartment-specific PKA substrate phosphorylation in hippocampal CA1 pyrami-dal neurons exhibits consistent dynamics not constrained by sensor limitations. a, Heat map of FLIM-AKAR binding fraction across every NREM to wake transition from an example experiment. b, Heat map of FLIM-AKARm binding fraction across every wake to NREM transition from an example experiment. c, Example trace of FLIM-AKARm binding fraction in medial prefrontal cortex (mPFC) in response to optical stimulation of photoactivatable adenylate cyclase (biPAC; stimulation parameters: wavelength=470nm, duration=100ms, power=128μW). Green region denotes wake. Blue dashed line denotes time of stimulation. d, Average traces of FLIM-AKARm binding fraction change during wake in response to blue light stimulation of biPAC at various durations (n=2 mice). Data show mean ± SEM across epochs. e-f, Example traces (e) and summary of decay constants (τ) from exponential fitting (f) of FLIM-AKARm binding fraction in the mPFC during baseline NREM bouts (n=5 mice) and following scopolamine injection (n=5 mice; NREM sleep vs. scopolamine injection, one-tailed Wilcox-on signed-rank). *: p<0.05.

**Extended Data 2.**
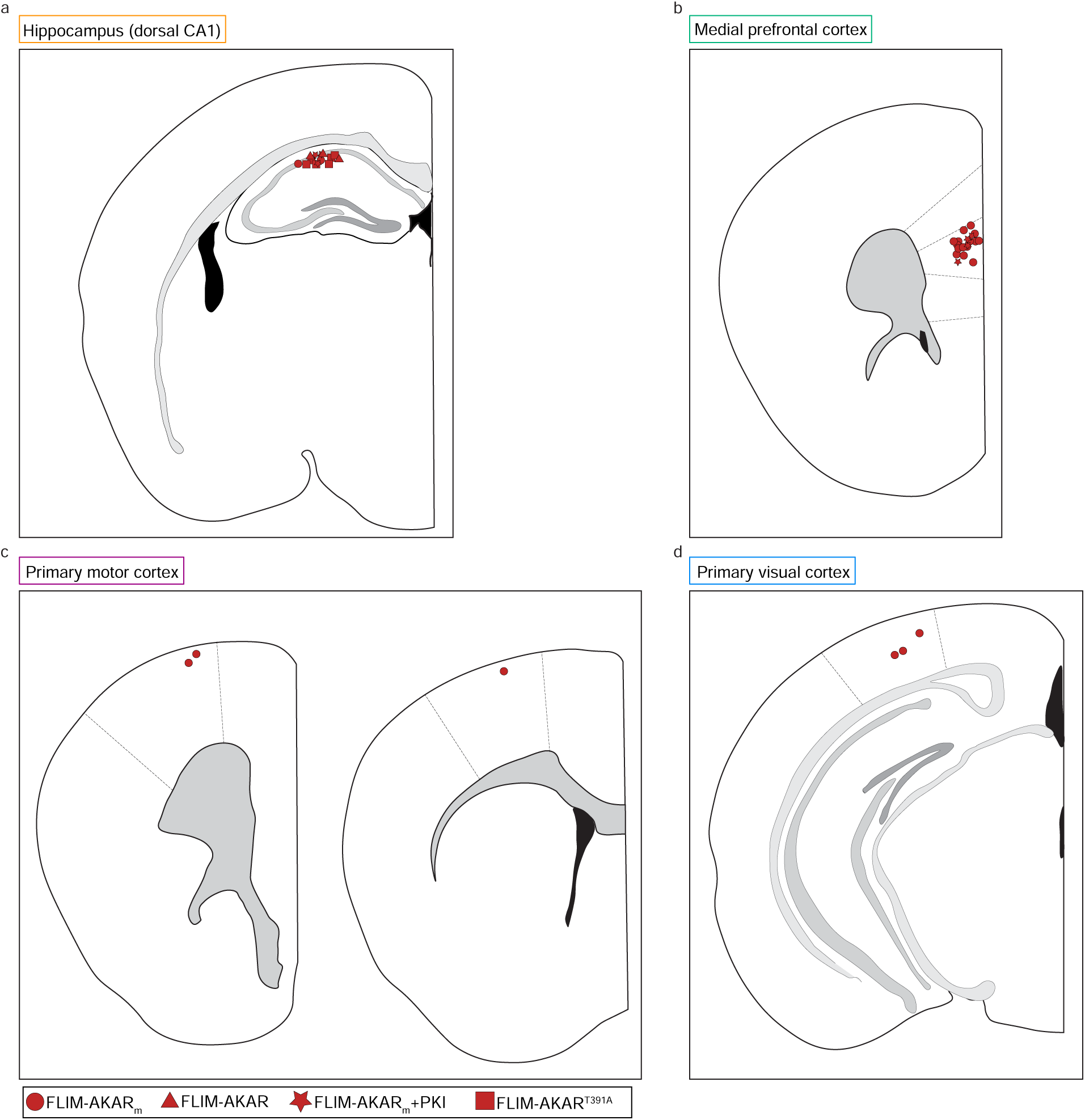
Summary of sensor expression locations across brain regions. a-d, Locations of expression of FLIM-AKAR, FLIM-AKARm, FLIM-AKARm+PKI, or FLIM-AKART391A in the (a) dorsal hippocampal CA1, (b) mPFC, (c) primary motor cortex (M1), and (d) primary visual cortex (V1).

**Extended Data 3.**
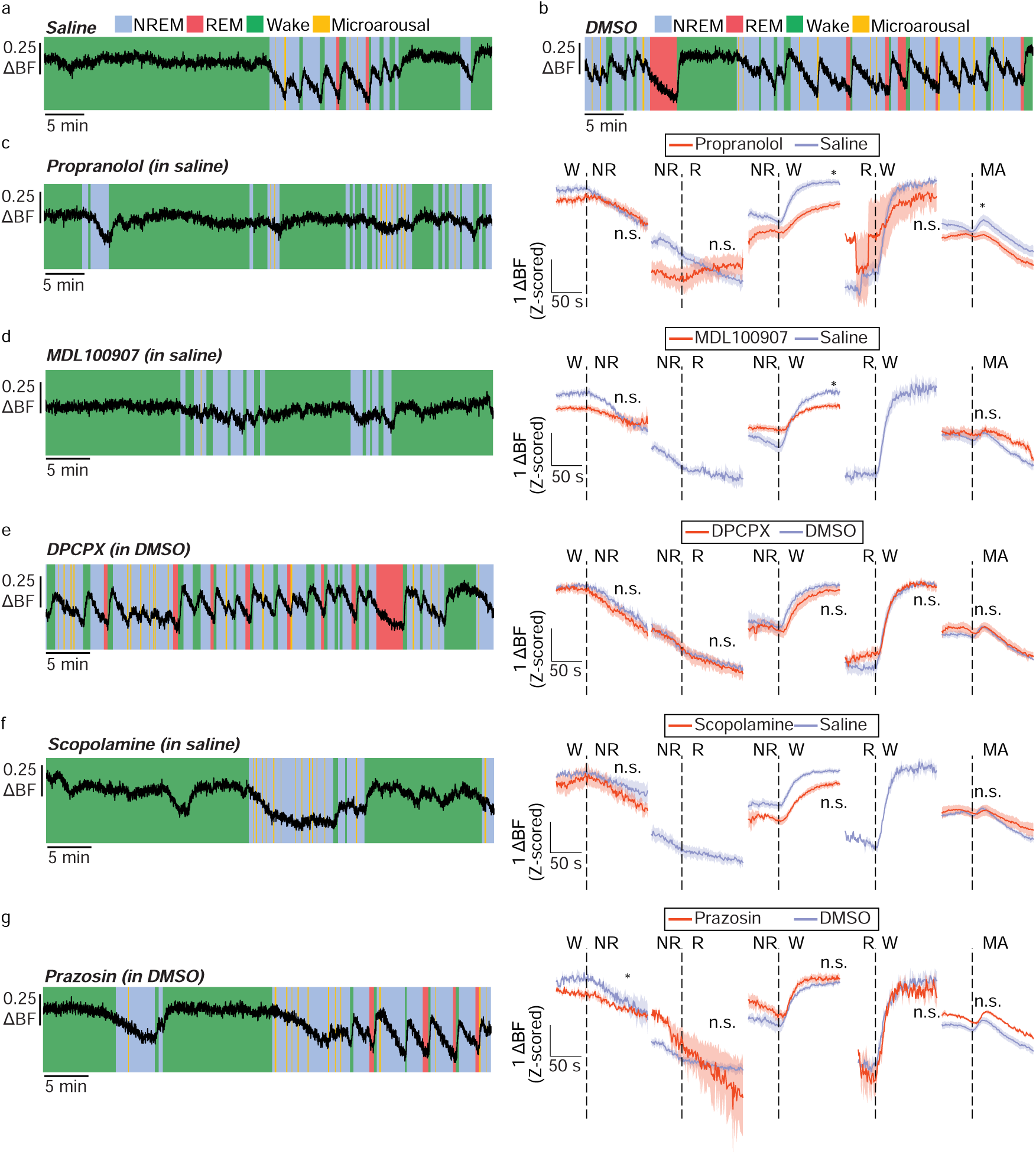
Effect of neuromodulator receptor signaling on sleep-wake-associated PKA-SP dynamics in the mPFC. Drug or vehicle was delivered every two hours during the light period. a-b, Example traces of FLIM-AKARm binding fraction following vehicle (saline or DMSO) injection. c-g, Example traces (left) and behavioral state transition-triggered averages (right, mean ± SEM) of vehicle or drug injections of (c) the β-adrenergic receptor antagonist propranolol (n=7 mice; see details in Extended Data Table 1) (d) the 5-HT2A receptor antagonist MDL100907 (n=4 mice; see details in Extended Data Table 1), (e) the adenosine receptor antagonist DPCPX (n=5 mice). (f) the muscarinic receptor antagonist scopolamine (n=5 mice; see details in Extended Data Table 1), and (g) the α-adrenergic receptor antagonist prazosin (n=5 mice; vehicle vs. drug; see details in Extended Data Table 1). *: p<0.05; n.s.: not significant, p>0.05.

**Extended Data 4.**
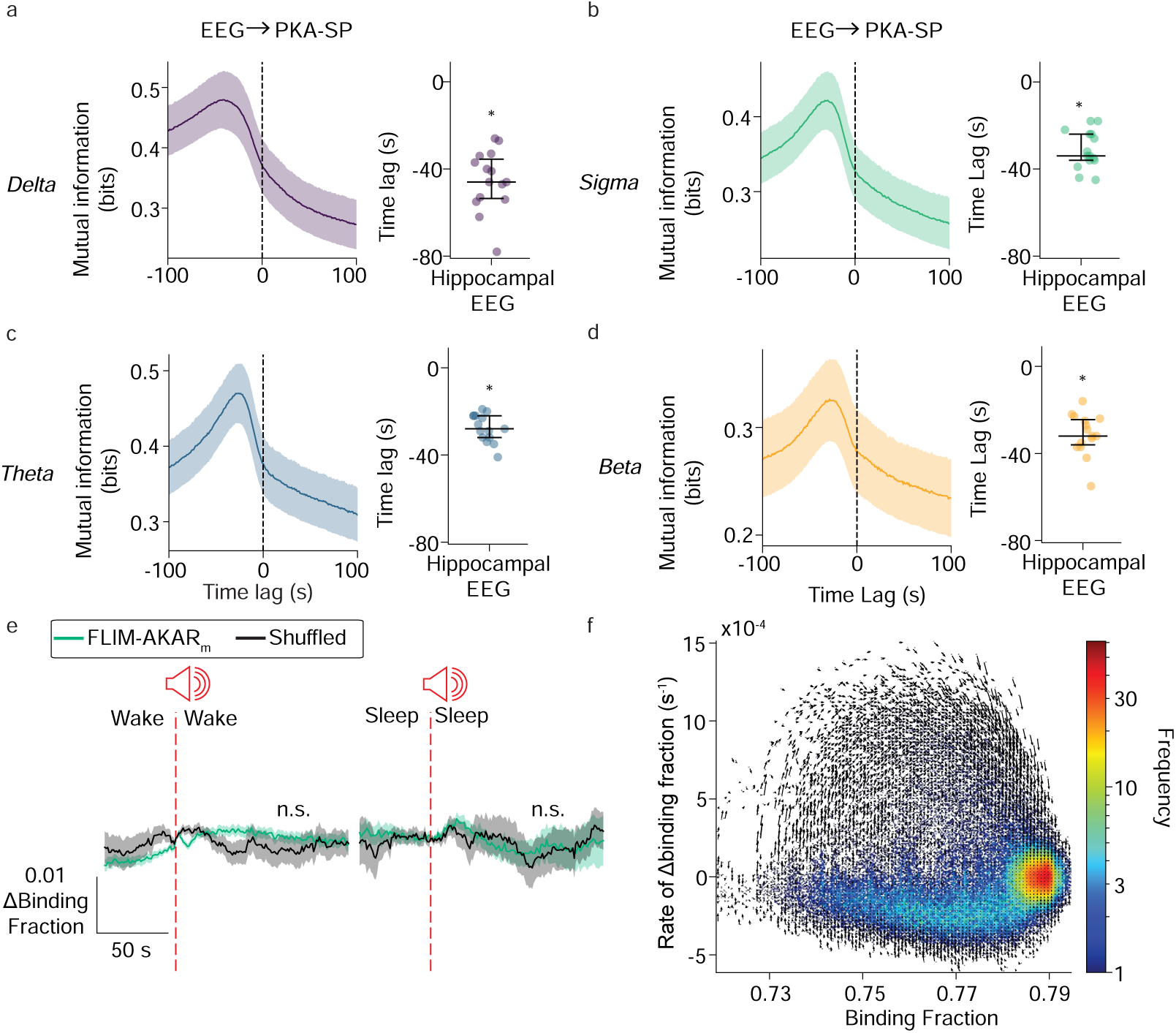
Membrane PKA-SP in the mPFC follows hippocampal EEG changes but not sound stimuli that do not induce state changes. a-d, Average traces (left; mean ± SEM) and peak lag (right; median with IQR) of mutual information between hippocampal EEG band powers and FLIM-AKARm binding fraction. Negative lag indicates EEG power precedes FLIM-AKARm binding fraction (n=15 mice; 1-sample Wilcoxon signed-rank). e, FLIM-AKARm binding fraction (n=3 mice) aligned to tone onset when the stimulus was delivered during a wake bout (left) or a sleep bout that did not result in a state change (right). Data show mean ± SEM. (FLIM-AKARm vs. shuffled; paired t-test). f, Two-dimensional histogram of binding fraction (x axis) vs. rate of change in binding fraction over time (y axis) for an example experiment. Arrows at each bin represent the average trajectory of the values in that bin over the next 30 seconds. *: p<0.05; n.s.: not significant, p>0.05.

**Extended Data 5.**
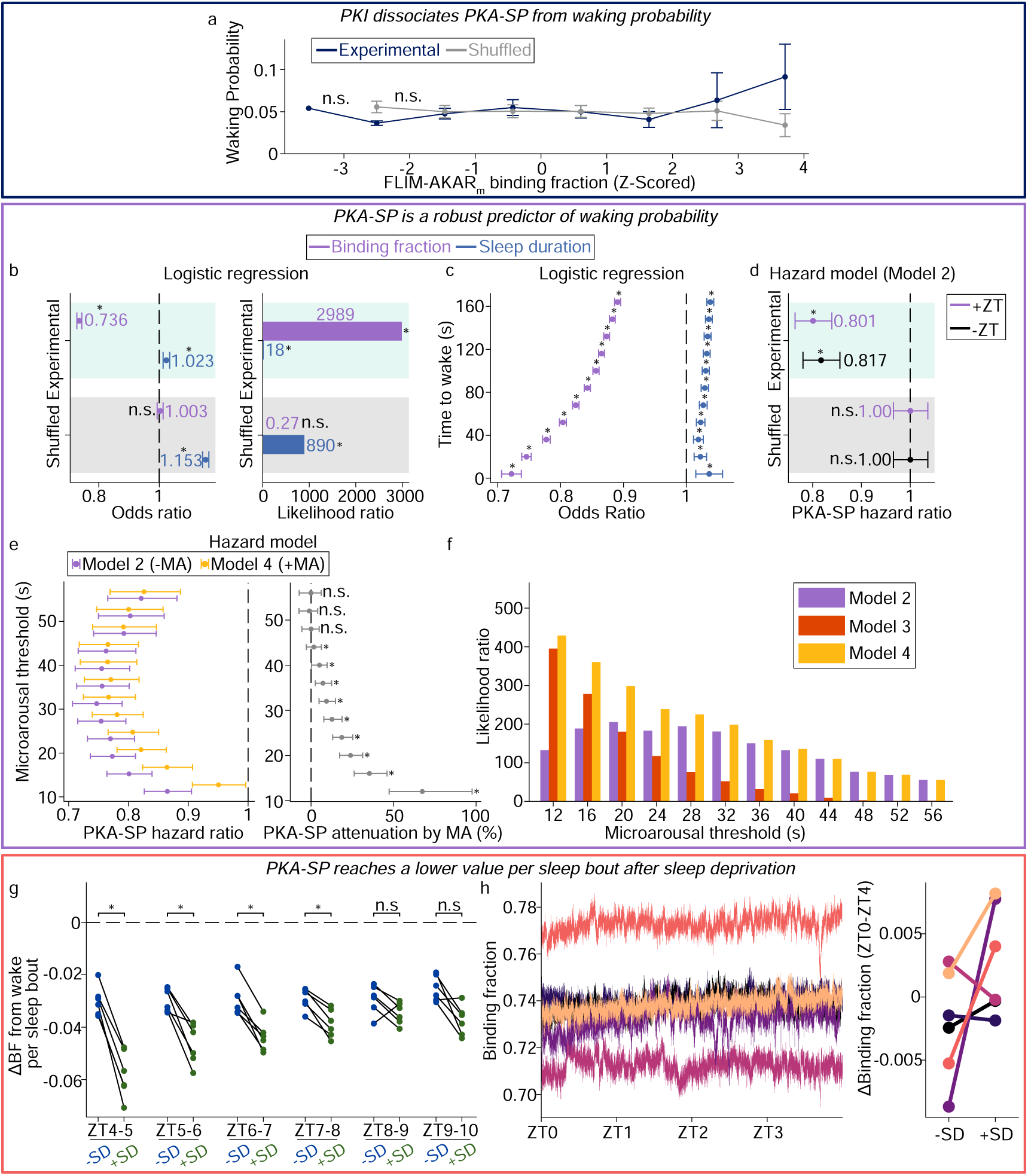
Membrane PKA-SP dynamics are a robust predictor of waking probability. a, Waking probability across Z-scored binding fraction of FLIM-AKARm in the mPFC in the presence of PKI. Data show mean ± SEM. (n=3 mice; mixed linear effects mode). b, Odds ratios (OR) ± 95% confidence intervals (CI) (left) and likelihood ratio (LR) (right) of FLIM-AKARm binding fraction and sleep duration from logistic regression models predicting waking probability in the next 16 seconds. The vertical dashed line at OR of 1 indicates no effect. (n=15 mice; OR: Wald test, LR: likelihood ratio test). c, Odds ratio of FLIM-AKARm binding fraction and sleep bout duration for predicting imminent waking in a range of subsequent time windows (4-160 seconds). Data shown with 95% CI. (n=15 mice; Wald test). d, Hazard ratio (HR) of FLIM-AKARm binding fraction with and without Zeitgeber Time (ZT) as an additional covariate (n=18 mice; shown with 95% CI; Wald test against null HR=1). The FLIM-AKARm binding fraction HR with ZT is reproduced from Fig. 5d for comparison. e, HR of FLIM-AKARm binding fraction (left; 95% CI) and attenuation of the log-HR when adding microarousal count as a covariate (right; point estimate with 95% bootstrap CI; one-sided permutation test) across microarousal length thresholds from 12 to 56 s (n=18 mice). f, Likelihood ratio assessing the hazard model contribution of Model 2, 3, and 4 for different microarousal duration thresholds (n=18 mice). g, Per-animal mean change in FLIM-AKARm binding fraction between average waking level and end of each sleep bout, in animals with and without sleep deprivation (SD). Microarousals were treated as wake (n=6 mice; −SD vs. +SD; Wilcoxon signed-rank). h, Example traces (left) and summary (right) of the change in FLIM-AKARm binding fraction between ZT0-ZT4, the sleep deprivation period or the circadian matched control (n=6 mice). *: p<0.05; n.s.: not significant, p>0.05.

**Extended Data 6.**
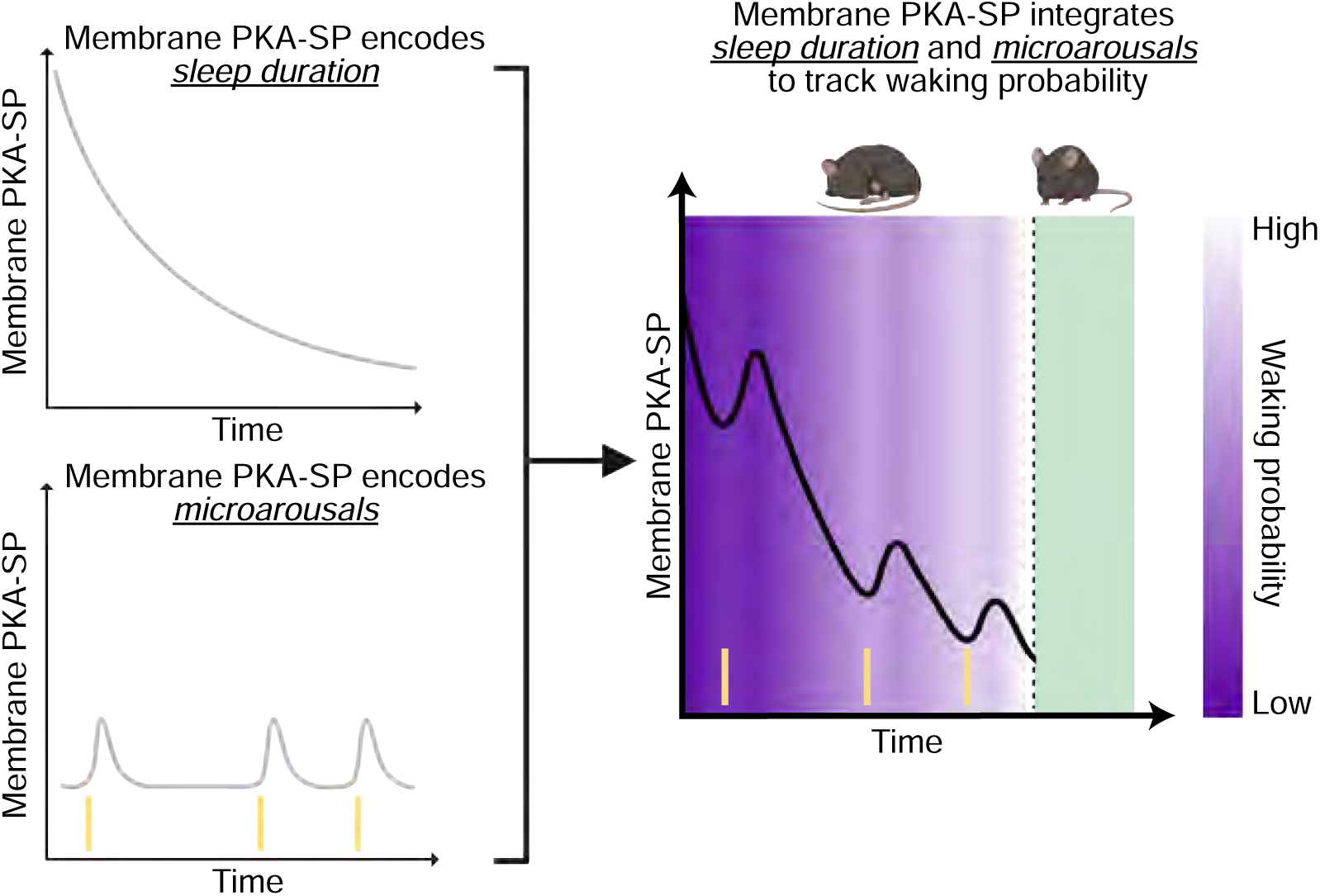
PKA-SP integrates sleep duration and sleep interruption to encode moment-to-moment waking probability within a sleep bout. Schematic illustrating our conclusion of how PKA-SP dynamics during sleep, via its dissipation kinetics and responses to microarousals, integrates sleep duration and sleep interruption to predict moment-to-moment waking probability.

